# Genomic characterization of a diazotrophic microbiota associated with maize aerial root mucilage

**DOI:** 10.1101/2020.04.27.064337

**Authors:** Shawn M. Higdon, Tania Pozzo, Nguyet Kong, Bihua Huang, Mai Lee Yang, Richard Jeannotte, C. Titus Brown, Alan B. Bennett, Bart C. Weimer

**Affiliations:** Department of Plant Sciences, University of California, Davis, California 95616; Department of Population Health and Reproduction, University of California, Davis, California 95616; 100K Pathogen Genome Project, University of California, Davis, California 95616

**Keywords:** Biological Nitrogen Fixation, Diazotrophic Bacteria, Mucilage Polysaccharide, Nitrogen Fixation Gene, *Zea mays*

## Abstract

A geographically isolated maize landrace cultivated on nitrogen-depleted fields without synthetic fertilizer in the Sierra Mixe region of Oaxaca, Mexico utilizes nitrogen derived from the atmosphere and develops an extensive network of mucilage-secreting aerial roots that harbors a diazotrophic microbiota. Targeting these diazotrophs, we selected nearly 600 microbes from a collection isolated from these plants and confirmed their ability to incorporate heavy nitrogen (^15^N_2_) metabolites *in vitro*. Sequencing their genomes and conducting comparative bioinformatic analyses showed that these genomes had substantial phylogenetic diversity. We examined each diazotroph genome for the presence of *nif* genes essential to nitrogen fixation (*nif*HDKENB) and carbohydrate utilization genes relevant to the mucilage polysaccharide digestion. These analyses identified diazotrophs that possessed canonical *nif* gene operons, as well as many other operon configurations with concomitant fixation and release of >700 different ^15^N labeled metabolites. We further demonstrated that many diazotrophs possessed alternative *nif* gene operons and confirmed their genomic potential to derive chemical energy from mucilage polysaccharide to fuel nitrogen fixation. These results confirm that some diazotrophic bacteria associated with Sierra Mixe maize were capable of incorporating atmospheric nitrogen into their small molecule extracellular metabolites through multiple *nif* gene configurations while others were able to fix nitrogen without the canonical (*nif*HDKENB) genes.

**Data Summary:** Genetic resources, including biological materials and nucleic acid sequences, were accessed under an Access and Benefit Sharing (ABS) Agreement between the Sierra Mixe community and the Mars Corporation, and with authorization from the Mexican government. An internationally recognized certificate of compliance has been issued by the Mexican government under the Nagoya Protocol for such activities (ABSCH-IRCC-MX-207343-3). Any party seeking access to the nucleic acid sequences underlying the analysis reported here is subject to the full terms and obligations of the ABS agreement and the authorization from the government of Mexico. Individuals wishing to access nucleic acid sequence data for scientific research activities should contact Mars Incorporated Chief Science Officer at.

## Introduction

Nitrogen is an essential macroelement for plant productivity that is often limiting to plant growth when the natural abundance of its bio-available forms is depleted in the environment. Exogenous nitrogen is currently provided for maize cultivation either through synthetic Haber-Bosch fertilizer produced at high environmental and economic cost (1), or from crop rotation with legumes that replenish field nitrogen levels by symbiotic association with diazotrophs, bacteria capable of biological nitrogen fixation (BNF) (2, 3). Because maize is a crop of immense agricultural importance, the establishment of conventional varieties capable of meeting their nitrogen demands through mutualistic associations with free-living diazotrophic bacteria would be of significant value to the goal of achieving global food security through sustainable intensification without relying on fertilization (4). One strategy for the discovery of useful maize diazotrophic plant-microbe associations involves exploring the microbiome of cultivated maize landraces near the center of the maize origin of domestication (5).

A recent report demonstrated that an indigenous landrace of maize found in Totontepec Villa de Morelos in the Sierra Mixe region of Mexico acquires 28-82% of its nitrogen from the air and exhibits an extensive system of aerial roots with heavy secretion of a mucilage composed of unique complex polysaccharides (6). Analysis of a public, low coverage shotgun metagenome sequences from the roots, stems, and aerial root mucilage revealed the aerial root mucilage microbiota to be enriched in taxa with many known species that are diazotrophic (6). In addition, the mucilage was the only plant tissue type to be enriched for homologs of the canonical nitrogen fixation genes (*nif*HDKENB), as previously proposed by Dos Santos et al., to be essential for a bacterium to be diazotrophic (6, 7). The demonstration that the Sierra Mixe mucilage harbors a diazotrophic microbial community, that it exhibits reduced taxonomic complexity, and the absence of soil from aerial root mucilage suggests that it could be a useful model system for elucidating associative mechanisms between free-living bacteria and cereal crops with mucilage-secreting aerial roots, such as maize.

Following investigations reported by Van Deynze et al. (6), we hypothesized that free-living diazotrophs from the aerial root mucilage microbiota utilize mucilage derived carbohydrates as an energy source for BNF. To address this, we cultured many bacteria by targeting diazotrophic bacteria specifically associated with Sierra Mixe maize. Subsequenctly, we characterized 588 microbial diazotrophic isolates to verify fixation and other traits using whole genome sequencing (WGS). Measuring the ability to incorporate heavy dinitrogen gas (^15^N_2_) into secreted metabolites with tandem mass spectrometry confirmed that the isolates were diazotrophic and produced a variety of compounds containing the label. Subsequent WGS analysis using comparative genomics with each diazotrophic isolate genome included assessing differences in nucleotide composition, assigning taxonomic classifications, and estimating percent recovery from the mucilage microbiome. To elucidate the genomic determinants for BNF by mucilage-derived diazotrophs, we examined their genomes for the presence of features related to mucilage polysaccharide utilization, the canonical *nif* genes based on the Dos Santos model (7) with the *Klebsiella pneumoniae* NIF regulon as the model framework, and known alternative *nif* genes. Our results indicate that the mucilage microbial isolates contained the capacity to utilize the mucilage complex polysaccharide and, surprisingly, that many of the diazotrophic isolates did not possess recognizable homology for known *nif* genes – yet were diazotrophic. These findings suggest the presence of novel mechanisms of nitrogen fixation by many phylogenomic groups of bacteria, several of which were not previously associated with this trait.

## Methods

### Bacterial isolation

Roots, stems and mucilage (200–500 μL) collected from different fields of the Sierra Mixe region in Mexico were spread on 1.5% BHI (BD, catalogue number 211059; Franklin Lakes, NJ, USA) or modified nitrogen-free M9 agar (BD) with and without a 1% (w/v) D-arabinose, galactose or xylose at pH 5, 5.8 or 7. Plant tissues were blended in 1×PBS prior to culturing on medium and the blender decontaminated with 10% bleach followed by 70% ethanol between samples (v/v). Cultures were incubated at 25°C or 37°C, aerobically and anaerobically, for up to 4 weeks. Once colonies appeared, they were sub-cultured on the same medium to ensure purity. Each organism was grown in BHI broth at the respective condition and resuspended in 5% non-fat dry milk and glycerol and stored cryogenically for further use.

### Biological Nitrogen Fixation Assay

To assay for Microbial ^15^N_2_ assimilation, isolates were first grown twice overnight in the respective growth condition prior to collection and washed twice with 0.9% (w/v) saline solution before re-suspension in Fahraeus medium containing 1% D-glucose at pH 5.8 to determine the nitrogen fixation capacity. Prior to the fixation assay, dissolved oxygen was removed from the medium by sparging with argon gas for 1.5 hours while stirring and a vacuum pump was used to remove any oxygen in the headspace. Each isolate (OD_600_ = 2; 2 mL) was added to an airtight 4 mL glass vial. Addition of the heavy atom was achieved by removing 20 mL of headspace gas and replacing it with 5 mL of either ^15^N_2_ or ^14^N_2_ nitrogen gas directly into the culture. The cultures were incubated at 37 °C anaerobically for 6 - 48 hours, depending on the growth rate and collected at the beginning of stationary phase for each culture. All experiments were done in triplicate.

### Microbial metabolite extraction and quantitation

Subsequent to growth the metabolites were extracted from cell pellets as described by Villas-Bôas (8). Bacterial cultures were transferred to 2 mL tubes and centrifuged at 14,800 rpm for 10 min at –9 °C. After collection of the cell pellet 500 μL of cold methanol (−20°C) was added before lysing the cells with bead beating (9, 10). After adding 0.4 g of 0.1 mm glass beads cells were lysed by two cycles of bead beating with 30 s per cycle, 1 min rest on ice between each cycle ^9,10^. The lysed samples were centrifuged at 14,800 rpm for 10 min at –9 °C after which 50 ml of each supernatant was transferred to LC vials for metabolite analysis. Samples were stored in -80 °C until analysis using LC/TOF-MS. In order to confirm the enrichment by ^15^N, a subset of residual pellets (50 mg of dried pellets), after metabolite extraction, were submitted to the UC Davis Stable Isotope Facility for Isotope Ratio Mass Spectrometry (IRMS) analysis (^15^N/^14^N ratio). ^15^N-labeled metabolite analysis was performed using LC-TOF G6230A (Agilent Technologies) instrument equipped with 1290 Infinity HPLC system. Chromatographic separation was performed on a Zorbax Eclipse XDB-C18 (2.1×15 mm, 1.8 µm) with a flow of 500 µL·min^-1^ and the following elution gradient: 0 min, 10 % B; 2.5 min, 80 % B; 4.0 min, 100 % B; 4.5 min, 100 % B; 5.0 min, 10 % B; 6.0 min, 10 %. Solvent A was water and solvent B was acetonitrile, both containing 0.1 % formic acid with a column temperature of 40 °C and an injection volume of 1-5 μL. This HPLC system was connected to an Agilent 6230 time-of-flight analyzer with an Agilent Jet Stream electrospray (ESI) interface operating in positive ion mode under the following conditions: capillary 3500 V, nebulizer 35 psi g, drying gas 8 L·min^-1^, gas temperature 350 °C, skimmer voltage 80 V, fragmentor voltage 135 V, octapole RF 750 V. The mass axis was calibrated using the mixture provided by the manufacturer in the m/z 50–1700 range. Acquisition rate was set to 1 spectrum per second (13,593 transients/spectrum). A reference solution provided continuous calibration using the following reference masses: 121.0509 and 922.0098 m/z. Accurate mass spectra from 70 to 1700 m/z were recorded and processed with MassHunter Workstation software (B.04.00). Statistical analysis was performed using GeneSpring-MassProfiler Pro (version 12.1) software from Agilent Technologies, and MetaboAnalyst (http://www.metaboanalyst.ca/) (11).

### Biomarkers of nitrogen-fixation

The basis of this approach is that as a microbe incorporates ^15^N by fixation, ^15^N will be used in the biosynthesis of small molecules and macromolecules, such as nucleic acids and proteins, shifting their masses of 1 unit per atom of nitrogen replaced. A given bacteria fixing nitrogen and exposed to ^15^N_2_ gas will have a very different spectrum compared to the same bacteria exposed to ^14^N_2_ only.

The mass spectrometry analysis of each extract generated an average spectrum per sample that contains thousands of masses. All the spectra were aligned and assembled in one data matrix using SpecAlign software. Using the data from all the isolates, we performed a statistical analysis (t-test, in MetaboAnalyst) (11) to determine the features (masses) that were significantly changing across isolates when controls and treated samples were compared. This approach allows us to identify biomarkers of nitrogen fixation that could be common to all the isolates, totally or partially (some isolates could have all the biomarkers identified, some others only a subset). More than 700 masses were significantly different using a q value (a p-value adjusted by False Discovery Rate (FDR); this statistical approach allows to correct for possible false positives) of 0.05 as threshold (q value ≤ 0.05 was determined to be significant). Masses with q≤0.05 and fold-change (intensity of given mass in 15N-treated samples vs intensity of the same mass in ^14^N-treated samples) of >1 were considered in the following calculations. Then for each isolate, the relative intensities (percentage of each peak raw intensity over total raw signal) for all the biomarkers were summed. Sums of the relative intensities for the biomarkers in control and treated samples, for a given isolate, were computed and ratio ^15^N/^14^N was calculated. Isolates with BNF ratios greater than or equal to 1 were considered as sufficient N_2_-fixers, where the sum of peak intensities under ^15^N_2_-enriched atmosphere was found to be equal to that of the unenriched control. Following this logic, isolates with BNF ratios greater than 1 were considered to be more efficient N_2_-fixers (i.e. higher ^15^N ratios indicated a higher detected abundance of ^15^N atom incorporation into N-containing biomarkers) while those with ratios lower than 1 were considered low-fixing.

### Bacterial whole genome sequencing

Each Sierra Mixe microbial isolate was recovered from cryogenic storage by streaking cells onto Luria-Bertani (LB) agar medium plates and incubating for one to two days at 28 °C. Single colonies were sub-cultured in liquid LB medium at 28 °C to an OD_600_ value of 0.7. Genomic DNA (gDNA) was extracted from the cell culture pellet of each isolate using the *Mo Bio* Ultraclean Microbial DNA extraction kit (QIAGEN, Inc). Sequencing libraries were subsequently constructed using the KAPA HyperPlus DNA library preparation kit (Roche, Inc) by following the instructions of the technical datasheet provided. A gDNA input of 100 ng was fragmented enzymatically for 9 minutes to achieve an average insert size of 450bp. The inserts were ligated to customized dual-indexed barcode adapters (Integrated DNA Technologies), and the library was size-selected by using KAPA Pure beads to carry out the kit’s dual-SPRI protocol to generate an average adapter-ligated gDNA insert molecule size of 600 bp. The size-selected libraries were then PCR amplified over a total of five cycles. Average library molecular size was determined using the DNA High Sensitivity Assay kit with the Agilent 2100 Bioanalyzer (Agilent Technologies). The Library was then used to generate paired end reads over 150 cycles at the UC Davis DNA Sequencing Technologies Core facility on the Illumina HiSeq 4000 system.

### Isolate Genome Sequence Analysis

The paired-end FASTQ files of each isolate library were quality trimmed using Trimmomatic 0.36 using the following settings: ILLUMINACLIP:TruSeq3-PE.fa:2:40:15; LEADING: 2; TRAILING:2; SLIDINGWINDOW:4:15; MINLEN:50 (12). The trimmed reads were subsequently assembled using MEGAhit 1.2 with default settings (13). Assembly metrics were obtained with the default settings of QUAST 4.1, the quality assessment tool for genome assemblies (14), and the output for each assembly is summarized in S2 Table. Genome binning analysis to assess the purity of each isolate genome was carried out using the program Metabat with the default settings (15). The number of bins generated by Metabat for each isolate genome are displayed in S2 Table. Values for genomic coverage were generated by aligning trimmed reads to the resulting assemblies with BWA followed by the use of the depth function from Samtools (16, 17). Code for the Snakemake workflow used to conduct the computational analysis is available at: (https://github.com/shigdon/snakemake_mucilage-isolates).

### Genome distance analysis and taxonomic classification

Whole genome assemblies were classified and compared using Sourmash 3.1.0 (18), which provides implementation of both the MinHash and Lowest Common Ancestor (LCA) algorithms to carry out whole genome comparisons and taxonomic classification of microbial isolates in a fast, efficient and lightweight computational fashion (18-20). The complete assembly files output from MEGAhit 1.2 for each isolate genome were used to generate MinHash signatures, also referred to as sketches, using the program Sourmash 3.1.0 (https://github.com/dib-lab/sourmash). The chosen k-mer size for each isolate genome’s MinHash signature was set to 31 (k-31). These sketches served as genomic fingerprint signatures that were used to carry out an all-by-all comparison at the whole-genome level by using the ‘compare’ function of Sourmash to calculate Jaccard Similarity Index (JSI) values for each pairwise comparison, which was output as a matrix in csv format. This csv file was then used to generate the all by all comparative matrix and associated dendrogram in Fig. 1 using the ComplexHeatmap package in R (21). For taxonomic assignment of total genome assemblies, the k-31 signatures were queried against a database of k-31 MinHash signatures that correspond to the curated microbial genomes within the Genome Taxonomy Database (GTDB) v89 using the ‘lca search’ command of Sourmash (available at: https://osf.io/wxf9z/). K-31 MinHash signatures were also generated using Sourmash for the genome bins of each isolate genome that were created using Metabat. The MinHash signature of each genome bin was classified using the ‘lca search’ function of Sourmash using the aforementioned prepared database. Results from bin classification using Sourmash are presented in S4 Table. Quantification of full taxonomies generated using Sourmash LCA classification data from isolate genome bins derived was visualized as a Heat Tree using MetacodeR 0.3.1 in R (22).Code used to generate, compare and classify MinHash genome sketches is included in the Snakemake workflow hosted at: (https://github.com/shigdon/snakemake_mucilage-isolates). Code used for analysis of Sourmash output and figure generation in R is available at: (https://github.com/shigdon/R-Mucilage-isolates-sourmash).

**Fig 1.**
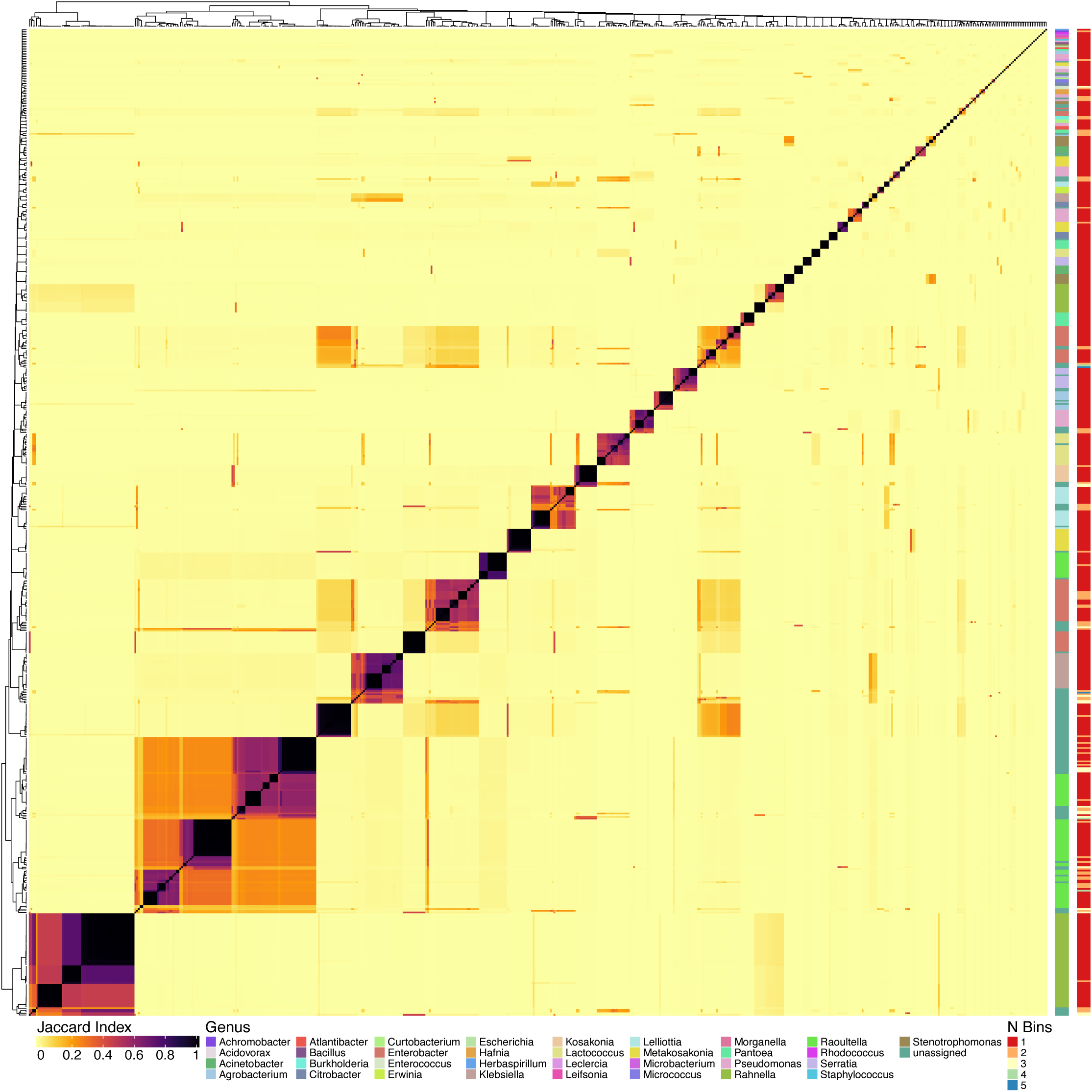
Comparative analysis of draft genome assemblies from Sierra Mixe bacterial isolates. All-by-all comparison of MinHash sketches of draft genome assemblies from 588 bacterial isolates using Sourmash (18). MinHash sketches of each draft genome assembly used in the comparison had a k-mer size of 31. Genus classification from MinHash sketches for each isolate genome is presented as a color-coded sidebar alongside the matrix. Results from genome binning analysis with Metabat (15) is included as a second color-coded sidebar. The Jaccard Index scale represents the Jaccard Similarity Index (JSI) value computed for each pairwise comparison of isolate genome MinHash sketches. Darker coloring indicates higher genome similarity and lighter coloring indicates lower similarity.

### Mucilage metagenome taxonomic classification

Paired end Illumina sequence data from Sierra Mixe aerial root mucilage metagenome sample OLMM00 was downloaded from Figshare (https://figshare.com/s/04997ae7f7d18b53174a#/articles/6615497) and analyzed to characterize the breadth of microbial diversity present within the mucilage environment. The shotgun metagenomic reads were quality filtered using Trimmomatic 0.36 and the surviving reads were separated into microbial and non-microbial fractions using the classify function of Kraken2 2.0.8_beta with the Refseq complete databases for Bacteria, Archaea, and Viruses (23, 24). The microbial component of OLMM00 classified with Kraken2 was subsequently visualized using the R package MetacodeR at the Phylum, Class, Order and Family levels, which is presented in Fig S1 (25). The relative abundance of each microbial taxon classified at the genus level was computed after performing Bayesian re-estimation of hits using Bracken2 (26) and normalization of read classifications for each taxon with the counts per million method using the R package Phyloseq (S6 Table) (27). Prior to analyzing the microbial community, the table of classified microbial taxa output by Bracken2 was filtered to remove taxa for which the number of classified reads was below 500, which resulted in a total of 609 unique genera identified within the OLMM00 metagenome (S7 Table). Source code for analysis and figure generation is available at: (https://github.com/shigdon/R-Mucilage-Metagenome).

### *Nif* and alternative *nif* gene mining

Protein coding sequences were predicted for each microbial isolate genome by using the corresponding MEGAhit-assembled contigs as input files for the prokaryotic genome annotation program Prokka 1.12 (28). The multi-FASTA amino acid files output for each isolate genome were scanned against profile hidden markov models (pHMMs) corresponding to *nif* genes of the *K. pneumoniae* NIF regulon using the ‘hmmscan’ function of HMMER 3.1b (29). These were acquired from the Pfam and TIGRFAM libraries of pHMMs (30, 31). HMM hits for each *nif* gene were stringently filtered in R using the dplyr package to retain query-subject hits that maintained model coverage greater than or equal to 75 % and a maximum e-value of 1e^-9^ (32). Visualization of *nif* gene profiles for all pure isolates depicted in Fig 3 was achieved using the Complex Heatmap package in R by clustering pure isolates based their relative MinHash distances and displaying counts of unique coding sequences that were found to match each *nif* HMM (21). TIGRFAMs used to scan for canonical *nif* genes of the *K. pneumoniae* NIF regulon included: TIGR01817, TIGR02938, TIGR02176, TIGR01287, TIGR01282, TIGR01286, TIGR01283, TIGR01285, TIGR01290, TIGR02000 TIGR03402, TIGR02660, TIGR02933 and TIGR01752. Pfams used to scan for *nif* gene mining included: PF04891.11 and PF03206.13. TIGRFAMs used to scan for alternative *nif* gene mining included: TIGR01860, TIGR02930, TIGR02932, TIGR01861, TIGR02929 and TIGR02931. The corresponding hmmscan results for alternative *nif* genes were filtered to retain query-model matches with maximum e-values of 1e^-06^ and 85 % minimum model coverage. Source code for bacterial genome mining analyses and figure generation is available at: (https://github.com/shigdon/R-Mucilage-isolates-nif) and (https://github.com/shigdon/R-alt-nif-analysis).

**Fig 2.**
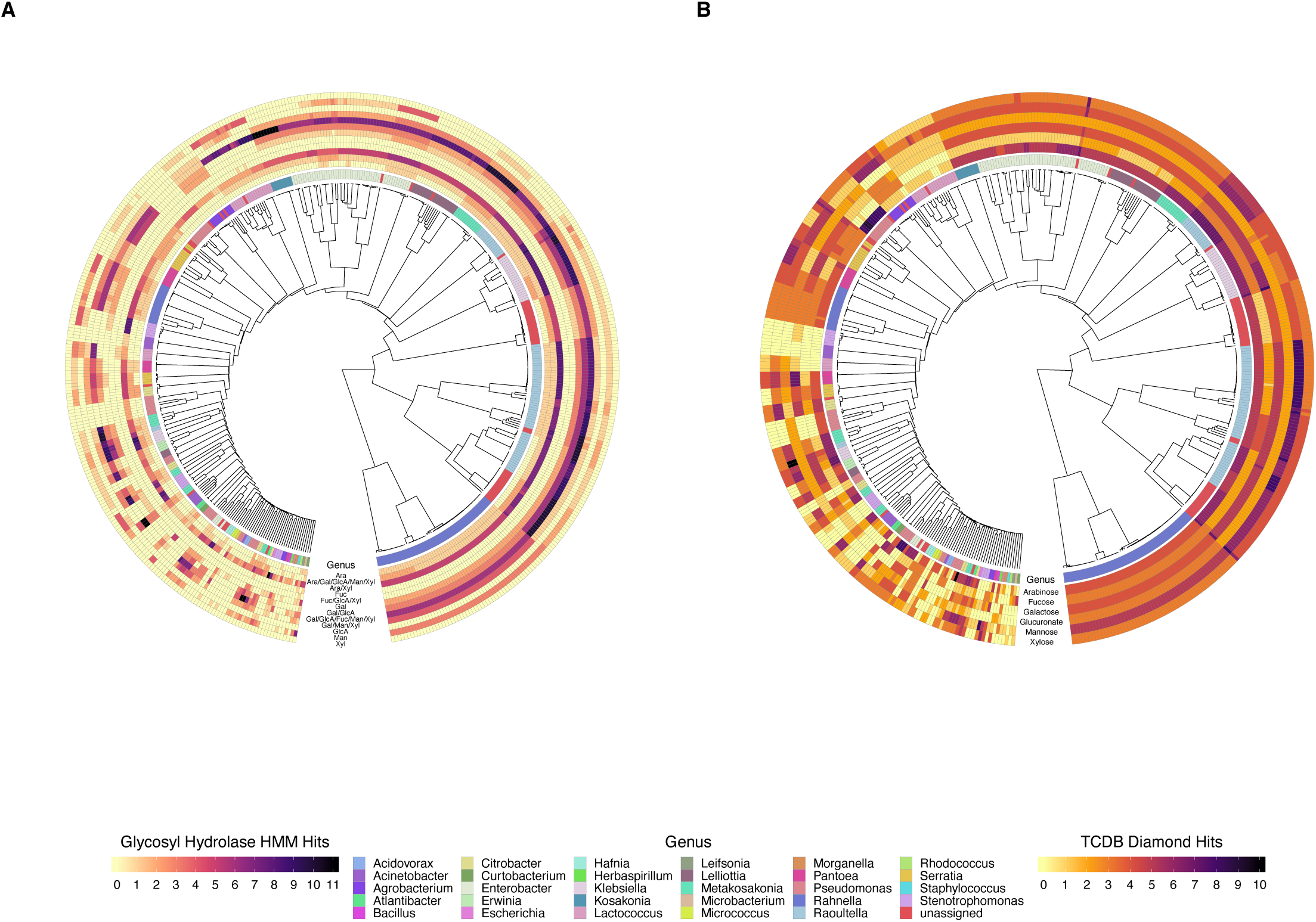
Glycosyl hydrolase and sugar transporter genome profiles of diazotrophic isolates. Analysis using dbCAN2 (33) was done to query total predicted coding sequences in each genome. Gene sequences encoding CAZymes and sugar transporters with substrate specificities that correspond to monosaccharide residues of the Sierra Mixe aerial root mucilage polysaccharide were selected from query results by generating a manually curated list of CAZy HMMs and TCDB accession IDs. Predicted gene sequence-HMM matching pairs were reported after filtering total hits from each genome to select all records with > 85% model coverage and an e-value ≤ 1e^-09^. A) Glycosyl Hydrolase family HMM hits with designated sugar residue specificities: Ara – Arabinose, Gal – Galactose, GlcA – Glucuronic Acid, Fuc – Fucose, Man – Mannose, Xyl – Xylose. B) Sugar Transporter HMM-Gene hits with designated sugar residue transporter activity.

**Fig 3.**
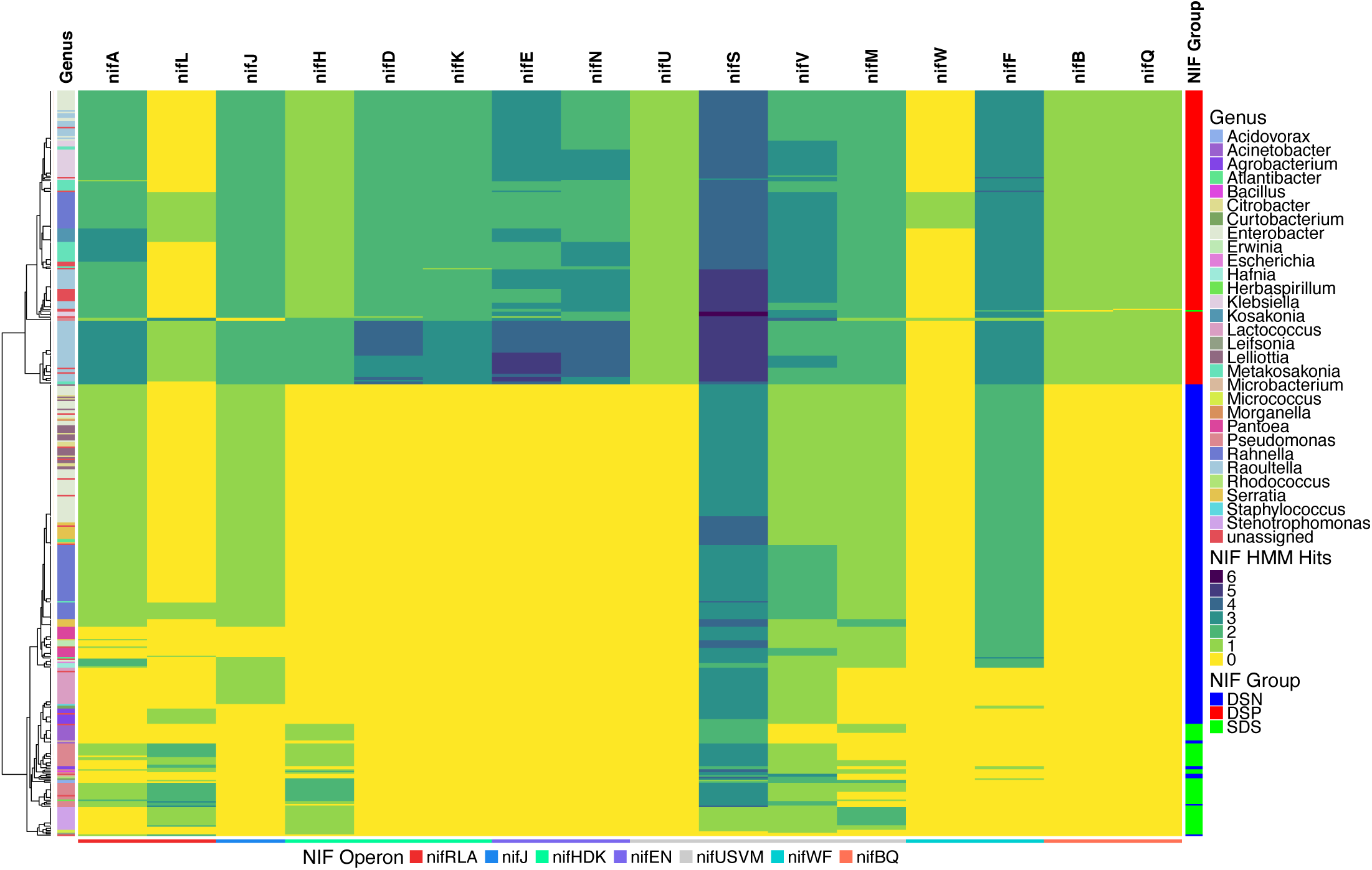
Canonical *nif* gene profiles of diazotrophic isolate genomes. Total predicted protein sequences of each pure isolate genome were queried against Hidden Markov Models (HMMs) for genes of the *K. pneumoniae* NIF regulon – including the six essential *nif* genes of the Dos Santos (DS) Model. Pure isolate genomes were clustered based on their relative MinHash genomic distances followed by heatmap visualization of their associated *nif* gene profiles. Three groups of pure diazotrophic isolates were formed based on the detected presence of homologous protein coding sequences to *nif*HDKENB: DS-Positive (DSP; red), Semi-DS (SDS; green) and DS-Negative (DSN; blue). Predicted amino acid sequence queries for each genome were considered as matches if *nif* gene HMM coverage was greater than or equal to 75% along with e-values ≤ 1e^-9^.

### CAZyme gene mining

The multi-FASTA amino acid files for each microbial isolate genome that were generated by Prokka were each used as input for the dbCAN2 analytical pipeline (33). This was achieved using a local installation of the source code for the dbCAN2 pipeline hosted on Github (https://github.com/linnabrown/run_dbcan). Output files in CSV format were read into R and filtered using the R packages within tidyverse 1.2.1 (34). Circular heatmap plots were made using the ggtree package (35). Source code for analysis and figure generation is available at: (https://github.com/shigdon/R-Mucilage-isolates-dbCAN2).

### Pan-genome Analysis

Genomic features predicted by Prokka (36) for each microbial genome included in the isolate sub-population study were aggregated in GenBank feature format and collectively used as input for pan-genome analysis using the program Roary 3.12.0 (37). Configuration for running the Roary microbial pan-genomic pipeline included use of the “-e” flag to generate a multi-FASTA alignment of core genes using PRANK and a minimum blastp identity value of 95 percent. To visualize the pan-genome of the isolate set presented in Fig 5C, the gene presence and absence output file, the associated dendrogram and an isolate-genus mapping file were uploaded to the Phandango web server (38). Source code for analysis and figure generation is available at: (https://github.com/shigdon/R-alt-nif-analysis).

**Fig 4.**
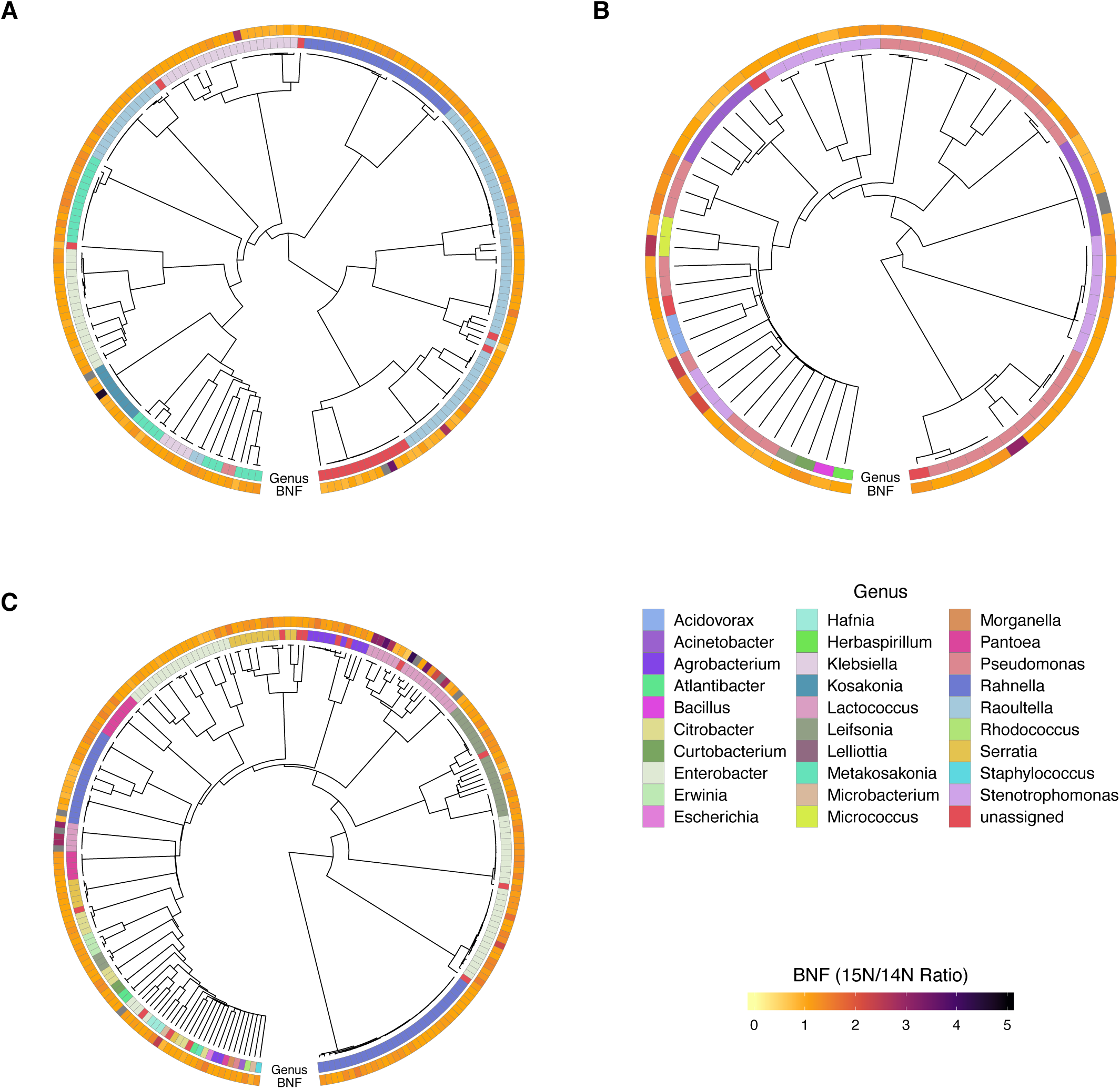
BNF ratios of mucilage diazotrophs from atmospheric ^15^N_2_ incorporation assay. As a means to connect each diazotroph’s *nif* gene profile with its corresponding BNF phenotype, BNF ratios are presented in heatmaps that accompany dendrograms clustered by MinHash genome distance under the context of the three NIF Groups determined from the genome mining analysis (Fig 3). Heatmap annotations indicate the ^15^N/^14^N ratios (BNF ratios) that represent the summation of peak intensities for all N-containing metabolites used as biomarkers in the assay. A) Dos-Santos Positive (DSP) isolates; B) Semi-Dos Santos (SDS) isolates; C) Dos-Santos Negative (DSN) isolates. Grey bars on the BNF ratio heatmap indicate values that were not determined.

**Fig 5.**
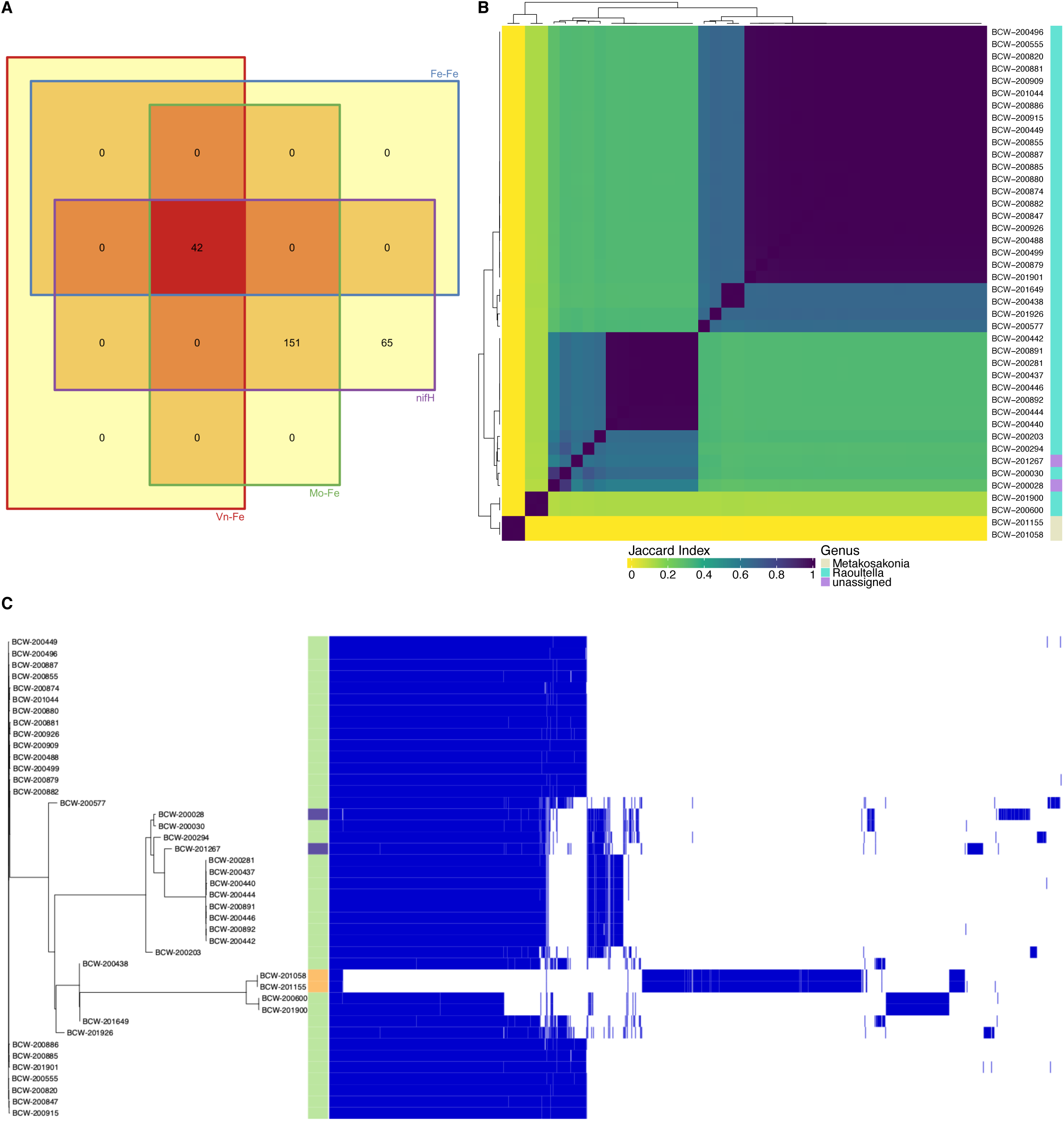
Mucilage bacterial isolates exhibit alternative nitrogenase genes. A) The presence of predicted protein sequences in diazotrophic isolate genomes was detected using TIGRFAM HMMs corresponding to the Fe-Fe and Vn-Fe alternative *nif* genes (*anf*D, *anf*K, *anf*G, *vnf*D, *vnf*K, *vnf*G) along with HMMs for *nif*HDK. Genomes with detected presence of the targeted genes were compared and quantified using a Venn Diagram to determine the list of diazotrophs with genes resembling Vn-Fe, Fe-Only, Mo-Fe Type nitrogenases and *nif*H. B) Genomes with alternative *nif* genes were compared using Sourmash and visualized as a composite matrix that included annotation of genus level classification. C) Pan-genome analysis of diazotrophs with alternative *nif* genes was conducted using Roary (37) and data for gene presence and absence was visualized using Phandango (73) along with genus classification data from Sourmash LCA. Orange annotations indicate genomes classified as *Metakosakonia*, green annotations indicate *Raoultella* isolates, and purple annotations indicate “unassigned” classification at the genus level.

## Results

### Diazotrophic isolates were confirmed by functional assay of ^15^N_2_ incorporation

We isolated putative diazotrophic bacteria in samples collected from Sierra Mixe maize plants grown using a nitrogen-deficient basal medium supplemented with sugars corresponding to the monosaccharide composition of aerial root mucilage (S1 Table). Culturing each isolate in N-deficient liquid media under an atmosphere containing ^15^N_2_ gas and measuring their ability to incorporate ^15^N atoms into small molecule metabolites (i.e. <1000 Da) by Time of Flight mass spectrometry confirmed that the isolates were diazotrophic and produced a large number of compounds with different masses and chemical structures. Summation of peak intensities for N-containing compounds common to enriched and control (compressed air) cultures enabled each isolate’s BNF capacity to be measured as a ratio of ^15^N/^14^N (BNF ratio). Overall, BNF ratios obtained for all pure isolates assayed ranged from 0.6 to 4.6 (Table S2). While most isolates exhibited moderate BNF ratios between 1 and 2, ∼5% of the isolates demonstrated N-fixation with BNF ratios >2 (Table 1). The observed BNF ratio variation among these confirmed diazotrophs prompted investigation of the underlying genomic determinants for BNF of each isolate.

**Table 1.** Summary of BNF Assay Results. Isolates were grouped using defined ranges of ^15^N/^14^N ratio values. ^15^N/^14^N ratios were computed by summing the peak intensities of all N-containing bio-markers common to both enriched and control cultures that had q-values less than or equal to 0.05 after analyzing metabolite data using Metaboanalyst (11).

| BNF Group | N Isolates | $^{15}\text{N}/^{14}\text{N}$ Ratio |
| --- | --- | --- |
| A | 4 | $x > 4$ |
| B | 10 | $4 > x > 3$ |
| C | 14 | $3 > x > 2$ |
| D | 461 | $2 > x > 1$ |
| E | 85 | $x < 1$ |
| F | 14 | Not determined |

### Whole genome analysis revealed significant phylogenetic diversity

The selected bacterial isolates were subjected to WGS and resulted in a collection of draft genome assemblies with fold coverages that ranged from 14 – 330X (S3 Table). Analysis of mucilage isolates revealed an unexpected range of diversity in nucleotide composition and taxonomy. All-by-all comparison of MinHash sketches for each isolate genome depicted the relative genomic distances of all pairings that verified the diversity of genomes (Fig 1). Complete taxonomic classification for each bacterial genome (S4 Table) at the maximum sketch size found 33 known bacterial taxa among the 472 isolate genomes, and 116 genomes that were unidentified (Fig 1). Possible explanations for unidentified isolates included lack of a database accession match or the presence of multiple bacterial genomes within a WGS MinHash sketch that triggered disagreement within the genomic classification structure of the lowest common ancestor algorithm (LCA).

To assess whether isolate genomes were pure or derived from a mixed culture that appeared pure during isolation, we used Metabat (15) to bin each WGS assembly and identify isolates comprised of multiple organisms. This resulted in 492 isolate genomes with single bins of single organism DNA sequences (Fig 1, S6 Table) – indicating pure cultures. WGS assemblies with 2, 3, 4 and 5 bins had frequencies of 72, 19, 3 and 2, respectively (Fig 1 and S3 Table), indicating that what appeared to be a single colony contained multiple organisms and that further WGS analysis was needed to deconvolute respective sequences. Reexamination of the deconvoluted genomes for taxonomic classification of each genome bin increased the resolution of microbial diversity and augmented the diversity of the taxa present and capable of fixing nitrogen (S5 Table).

Visualization of the classified genome bins indicated that the selected isolates were primarily comprised of Proteobacteria, a substantial number of Firmicutes, and relatively few Actinobacteria (S1 Fig). While deconvoluted genomes largely classified as Gammaproteobacteria, relatively few deconvoluted genomes were classified to the Alphaproteobacteria or Betaproteobacteria classes. Congruent with the findings of Carvalho et al., several deconvoluted genomes from our study were classified as *Burkholderia*, along with other Betaproteobacteria that included *Achromobacter, Acidovorax* and *Herbaspirillum* (39). However, deconvoluted genomes classified as *Enterobacter, Klebsiella, Metakosakonia, Rahnella, Raoultella*, and *Pseudomonas* were among the most abundant in the mixed cutures. Membership of deconvoluted genomes classified to Firmicutes included a substantial number of *Lactococcus* and several were identified as *Enterococcus* and *Bacillus*. Included in the few Actinobacteria genomes sequenced, deconvoluted genome analysis found *Curtobacterium, Leifsonia, Microbacterium, Micrococcus* and *Rhodococcus* as well.

Comparison of the deconvoluted genomes and pure genomes for taxonomic content with the OLMM00 mucilage metagenome reported by Van Deynze et al. (6) indicated that the culturing strategy enriched the isolates that fixed nitrogen and obtained a small fraction of the possible mucilage microbiome reported from the low sequence coverage metagenome. Using 609 genera identified in OLMM00 as a benchmark for bacterial diversity (S7 Table), the unique genera classified among isolate WGS assemblies comprised ∼5% of genera in the mucilage microbiome. In addition, analysis of OLMM00 metagenome provided further insight to the phylogenetic diversity of mucilage microbiota associated with this landrace (S8 Table). Proteobacteria, Bacteroidetes, Actinobacteria and Firmicutes were the most abundant phyla in the mucilage microbiome (S2 Fig, S8 Table). However, confirmation of multiple organisms contributing to mixed cultures (i.e. composite genomes) limited our ability to attribute observed BNF phenotypes to a distinct organism within co-cultured isolates. This observation prompted genomic profiling of each pure isolate genome for carbohydrate utilization and *nif* features to address the hypothesis that mucilage diazotrophs derive energy from mucilage polysaccharide to fuel BNF.

### Diazotrophic isolates possessed CAZymes and sugar transporters relevant for mucilage digestion

Examining isolate genomes for glycosyl hydrolase (GH) genes relevant to the composition of aerial root mucilage polysaccharide (6, 40) was done using Hidden Markov Models (HMMs) of GH families in the Carbohydrate Active Enzymes (CAZy) database (S9 Table) (41). This analysis revealed that the pure culture diazotrophs contained genes supporting the genomic potential to degrade and derive energy from mucilage polysaccharides. Targeting GHs with arabinofuranosidase, fucosidase, galactosidase, glucuronidase, mannosidase and xylosidase activities revealed that diazotrophic genomes with small differences in genome diversity contained similar GH profiles spanning 12 functional GH groups (Fig 2A). Comparison of GH groups conferring arabinofuranosidase and/or xylosidase activities demonstrated that the more promiscuous ‘Ara/Xyl’ GH group had the highest abundance with increased genome copy number for the majority of classified genomes. GH groups with exclusive galactose or mannose substrate specificities were also abundant in the isolates examined, where the sum of the isolates with genes in these GH groups was determined to be 366 of the 492 genomes (S10 Table). In contrast to the plethora of genomes found to possess pentose and/or hexose cleaving GHs, those with strict glucuronate and fucose specificities were far less abundant in the pure cultures. Interestingly, most genomes possessed genes in GH groups with promiscuous substrate specificities that encompassed the complete range of mucilage polysaccharide compositional diversity across five different GH families (GH1, GH2, GH31, GH4, GH30).

In addition to generating GH profiles, querying genomes for the presence of sugar transport genes relevant to monosaccharides that contribute to mucilage polysaccharide structure revealed that isolated diazotrophs possess the machinery necessary for transport of mucilage-derived monosaccharides obtained from the digestion of mucilage, indicating that the initiating step of catabolism was present in the genome (Fig 2B). Utilizing a list of mucilage relevant accessions (S11 Table) from the Transporter Classification Database (TCDB) (42), we generated sugar transport profiles for each genome. Summarizing genome counts by genus level classification demonstrated that those classified to the most common Gammaproteobacteria exhibited sugar transporters for all six monosaccharide moieties derived from the mucilage polysaccharide (S12 Table). Additionally, isolates of the most commonly classified genera possessed multiple genes and/or mechanisms for transport of each monosaccharide type in mucilage. Genomes assigned to less abundant genera tended to exhibit higher variation in sugar transporter profiles, where the absence of known carbohydrate transport systems corresponding to some, but not all components of mucilage polysaccharide was observed. This observation may explain how the culturing strategy resulted in reflecting abundant members of the mucilage microbiome.

### Diazotrophic isolates displayed genomic variation in canonical *nif* gene features

The genetic basis for BNF was established following more than 100 years of research, where numerous *nif* genes have been implicated as contributing factors to the phenotype with various operon configurations (43). We investigated the genomic mechanism for the diazotrophic phenotype (i.e. BNF) by examining the predicted coding sequences using HMMs for the six *nif* genes of the Dos Santos model (7) within the context of seven genetic operons comprising the *K. pneumoniae* NIF regulon, which included: 1) the operon of *nif* genes involved in regulation of the *nif* pathway, *nif*RLA; 2) the catalytic operon, *nif*HDK; 3) operons involved in formation of the functional Fe-Mo protein, *nif*EN and *nif*BQ; 4) an operon of genes involved in assembly of the functional enzyme complex, *nif*USVM; and 4) operons conferring genes associated with mediating electron transfer, *nif*J and *nif*WF (44, 45). Results from this extensive analysis generated *nif* gene profiles and revealed three distinct groups of diazotrophic isolates (NIF groups) based on *nif* gene content and variation in structure (Fig 3). NIF groups included a subset of 193 genomes positive for the presence of homologous protein-coding sequences to HMMs for all *nif* genes in the Dos Santos model (DS-positive, DSP), a smaller subset of 66 isolates with *nif* gene profiles reflecting a semi-complete set of Dos Santos model *nif* genes (Semi-DS, SDS) and a subset of 233 isolates that completely lacked genes with HMM homology for all Dos Santos model *nif* genes (DS-negative, DSN), yet phenotypically displayed diazotrophy.

Although each NIF group included genomes classified from a range of bacterial genera, each group also included diazotrophs with an “unassigned” taxonomic classification (S6 Table). However, DSP genomes positively classified to known genera were comprised entirely of Gammaproteobacteria assigned to *Enterobacter, Klebsiella, Kosakonia, Metakosakonia, Pseudomonas, Rahnella* or *Raoultella*. SDS isolates had much higher taxonomic diversity, where SDS group membership was attributed by pure isolates from Actinobacteria, Firmicutes and Proteobacteria. Within these three phyla, SDS isolate genera included *Acidovorax, Acinetobacter, Bacillus, Curtobacterium, Herbaspirillum, Leifonia, Micrococcus, Pseudomonas* and *Stenotrophomonas*. In a fashion similar to SDS isolates, DSN genomes were also composed of Actinobacteria, Firmicutes and Proteobacteria. Interestingly, DSN genomes displayed the highest taxonomic diversity among the three NIF groups by including *Acinetobacter, Agrobacterium, Atlantibacter, Citrobacter, Curtobacterium, Enterobacter, Erwinia, Escherichia, Hafnia, Lactococcus, Lelliottia, Metakosakonia, Microbacterium, Morganella, Pantoea, Pseudomonas, Rahnella, Rhodococcus, Serratia* and *Staphylococcus*. While genomes classified as *Enterobacter, Metakosakonia* and *Rahnella* were found in both the DSP and DSN groups, *Pseudomonas* genomes were present in all three NIF groups. In addition to *Pseudomonas*, commonalities between genera identified within the SDS and DSN groups included membership to *Acinetobacter* and *Curtobacterium*.

Every genome from a diazotroph in the DSP group possessed homologous protein coding regions to *nif* genes in the *K. pneumoniae* NIF regulon (Fig 3, S3 Fig). Importantly, diazotrophs in this group possessed homologs to the six *nif* genes of the Dos Santos Model and exhibited BNF ratios that confirmed their ability to fix atmospheric nitrogen. The majority of diazotrophs in the DSP group had moderate BNF ratio values within the inclusive range of 1 to 2, and four isolates exhibited capacity ratios > 2 (Fig 4A). While the 21 *Rahnella* genomes were the only subset found to possess homologs for all 16 *nif* genes investigated, the remaining 172 genomes lacked homologs to either the *nif*J, *nif*L, *nif*Q or *nif*W genes in variable degrees and/or combinations. However, these diazotrophs exhibited nearly identical *nif* gene profile compositions with the exception of slight variations in gene copy number. In the case of DSP isolates classified as *Enterobacteriaceae*, distinguished clades of *Enterobacter* and *Klebsiella* genomes each lacked homologous genes to *nif*L and *nif*W while clades of *Pseudomonas* and most *Rahnella* genomes were the only diazotrophs with homologs for the *nif*W gene. With respect to the *nif*H gene encoding the dinitrogenase reductase protein, 150 genomes in the DSP catagory had single copy homologs and 43 exhibited the presence of two copies. Overall, the *nif* gene content and BNF ratios of diazotrophs in the DSP group demonstrated that many mucilage diazotrophs adhered to the *K. pneumoniae* NIF regulon and Dos Santos models to conduct BNF.

All 66 members of the SDS group contained homologs to at least one, but not all, of the essential *nif* genes in the Dos Santos model (Fig 3, S4 Fig) and fixed nitrogen with BNF ratios similar to diazotrophs in the DSP group (Fig 4B). In a similar fashion to the DSP isolates, all SDS isolates were found to possess homologs for at least one copy of the *nif*H gene but interestingly two copies were detected in 15 diazotrophs of the SDS group. Genes homologous to dinitrogenase component I, *nif*D and *nif*K, were only found in a single isolate of the SDS group. Regarding the three *nif* genes involved with biosynthesis of FeMoCo, only a single SDS isolate (BCW-200147) possessed homologs to *nif*E and *nif*N, and genes matching the HMM for *nif*B were not detected in any SDS diazotroph genomes. Beyond Dos Santos’ model of essential *nif* genes, many SDS isolates possessed homologs for several genes in the *nif*RLA and *nif*USVM operons of *K. pneumoniae*, but genes involved with electron transfer (*nif*F and *nif*J) were not detected among the majority of isolates in this group. Despite lacking the complete set of *nif* genes in the Dos Santos Model, BNF ratios for isolates in this group ranged from 0.8 to 3.0. Taken together, *nif* gene analysis combined with the diazotrophic phenotype (i.e. BNF ratios) in the SDS group revealed that many mucilage isolates exhibited BNF activity without the presence of any essential *nif* genes defined by the Dos Santos model, suggesting that a novel mechanism of diazotrophy may be expressed in the microbiome of this landrace.

Contrary to the DSP and SDS NIF groups, the 233 diazotrophs in the DSN group completely lacked the presence of homologous protein coding sequences for all *nif* genes in the Dos Santos model (Fig 3, S5 Fig) and exhibited BNF ratios that rivaled those of diazotrophs in the other NIF groups containing gene matches to HMMs for all or part of the *nif* genes in the Dos Santos model (Fig 4C). Members of the DSN group lacked homologs for many *nif* genes constituting the NIF regulon of *K. pneumoniae*, and nearly all of them possessed coding sequences resembling genes of the *nif*USVM operon. While many DSN genomes encoded homologous genes to the BNF regulatory protein *nif*A, members of this group contained gene sequences that matched the *nif*L HMM to a much lesser extent. Contrary to the observed *nif* gene profiles of diazotrophs in the SDS group, observed trends for DSN genomes included presence of homologous sequenecs to the *nif*F and *nif*J genes involved with electron transfer. Similar to observations made with the other two NIF Groups, 188 DSN diazotophs exhibited BNF ratios between 1 and 2. Surprisingly, among all three NIF Groups, the DSN group presented the largest number of diazotrophs with BNF ratio values > 2. Collectively, *nif* gene profiles of DSN genomes and their corresponding BNF assay results demonstrated that these diazotrophs were capable of BNF without employment of any *nif* genes in the Dos Santos model and only a subset of the *K. pneumoniae* NIF regulon.

### Alternative *nif* genes were detected in isolates with substantial genome variation

Following queries for canonical *nif* genes of the Dos Santos model, we investigated whether the bacterial genomes encoded *nif* genes for known alternative nitrogenase systems that either strictly utilize iron (*anf*) or incorporate vanadium in place of molybdenum (*vnf*) as metal co-factors of dinitrogenase (43). Utilizing TIGRFAMs for the *anf*D, *anf*K, *anf*G, *vnf*D, *vnf*K *and vnf*G nitrogenase genes along with those of the Mo-Fe type nitrogenase (*nif*HDK), HMM analysis of predicted protein sequences from each genome revealed a small subset of diazotrophs with alternative *nif* genes. This resulted in the identification of 42 genomes with coding sequences that matched all nine *nif* HMMs (Fig 5A). Investigation of these *nif* genes also confirmed 146 diazotrophs in possession of the *nif*HDK operon without genes matching alternative nitrogenase HMMs, and 63 that had genes matching only the HMM model for *nif*H. Investigating the genomes with alternative *nif* genes revealed that each was previously assigned to the DSP group. This observation warranted further investigation of genomic similarities and differences between the 42 genomes with alternative *nif* genes.

WGS comparison of diazotrophs that contained alternative *nif* genes uncovered substantial phylogenomic diversity within the group. Computation of genomic distances between the 42 previously identified genomes revealed 12 distinct groupings of highly similar diazotrophs with JSI of nearly 1 (Fig 5B). Cross-referencing previously generated taxonomy for these alternative *nif-*possessing diazotrophs revealed two genera classifications. Among these taxonomic assignments, 38 isolates were classified to be *Raoultella*, 2 isolates were classified as *Metakosakonia*, and 2 were classified as *Enterobacteriaceae*. This indicated that the majority of diazotrophs with homologs to alternative *nif* genes had genomes with significant nucleotide similarity to reference genomes in the Genome Taxonomy Database (GTDB) classified as *Raoultella* (46). Interestingly, diazotrophs classified as *Raoultella* exhibited broad genomic diversity and formed multiple taxonomic clusters, with the two “unassigned” genomes interspersed among them, suggesting that they are near relatives of *Raoultella*. Comparison of the JSI values between genomes classified as *Raoultella* presented values ranging from 0.1 to 1. Additionally, the two *Metakosakonia* genomes presented strong dissimilarity to the other 40 isolates with JSI values close to zero for each pairing. These observations indicated large variation in genome composition for this subset of isolated diazotrophs and prompted subsequent exploration of the pan-genome among isolates that lack classical *nif* genome construction yet fix nitrogen.

Observed differences in nucleotide composition among genomes with alternative *nif* genes were expanded by elucidating the pan-genome for this group of diazotrophs. Annotated protein coding features of each genome served as inputs for pan-genome analysis to determine the core genome among diazotrophic isolates with alternative *nif* genes (37). Pan-genome analysis revealed a narrow core genome comprised by 285 of the 15,353 genes provided as input (S13 Table) with 3,374 soft core genes, 2,532 shell genes and 9,162 cloud genes occurring within 95 – 99, 15 – 95, and 0 – 15 % of diazotrophic genomes, respectively. Genome clustering based on the presence and absence of annotated genomic features (Fig 5C) was highly similar to that observed using MinHash, where the isolate groupings of the phylogenetic tree generated using the pan-genome corresponded with clades determined using genome distance differences (Fig 5B). Although taxonomic annotation of diazotrophs comprising the pan-genome suggested many distinct groups of *Raoultella* genomes (annotated in green), interspersion of the two “unassigned” genomes with small blocks of unique coding features (annotated in purple) among the defined clades of *Raoultella* corroborated findings from the MinHash analysis with blocks of core genes. Visualization of the pan-genome revealed the *Metakosakonia* clade (annotated in orange) of two diazotrophs (BCW201058 and BCW201155) as a near relative to the duo of distinguished *Raoultella* genomes (BCW200600 and BCW201900), which confirmed findings from the genome distance analysis. Furthermore, these four genomes possessed large blocks of features absent from the other 38 genomes in the group.

## Discussion

### Diazotrophic isolates represented a small fraction of the mucilage microbiome

The strategy to isolate diazotrophs focused on simulating the native environment of aerial root mucilage (anaerobic/microaerophilic, pH and temperature) in combination with nitrogen deprivation. This enabled providing various carbon sources associated with the mucilage polysaccharide to force expression of the metabolic traits that are likely associated with growth and survival on maize during *in vitro* isolation and selection (S1 Table). This was based on the two-component hypothesis that diazotrophs of the resident microbiota incorporate atmospheric nitrogen into various compounds via BNF, which is biologically powered by ATP when utilizing sugars derived from mucilage polysaccharides to fuel the energy needs of the energetically expensive transformation. Successful generation of a large isolate collection from mucilage with this strategy set the stage for further investigations to confirm the putative diazotrophic isolates. In response, this study established an *in vitro* functional metabolomic assay to quantify each isolate’s ability to incorporate heavy nitrogen into various extracellular metabolites, which both confirmed the diazotrophic nature of isolates in this collection and verified the efficacy of the strategy to recover diazotrophs (Table 1, Fig 4, S2 Table).

WGS of nearly 600 diazotrophic isolates provided a means to assess the taxonomic diversity of the isolate collection relative to that of the mucilage microbiome. Concerns of isolate misclassification were avoided by using whole genome analysis and composition to assign taxonomy for diazotrophic genomes rather than a conserved marker gene with higher sequence conservation (47, 48). Utilizing Kraken to classify genera derived from normalized read counts (49) of the previously reported OLMM00 mucilage metagenome (6) (S7 Table, S8 Table) identified 609 genera, of which the diazotrophic genome collection had 29 in common (S5 Table). This revealed ∼5% of the bacterial diversity from the aerial root mucilage microbiota is contained within the isolate collection and demonstrated that the cultured subpopulation had 25% of the top 20 most abundant known genera in the OLMM00 metagenome. Although many diazotroph genomes were “unassigned” taxonomically, which highlights the potential novelty of many bacteria in this isolate collection, metagenome sequencing of mucilage samples at a higher depth and re-classification of isolate genomes following expansion of microbial WGS databases should be achieved in the future to verify these results.

Comparing taxa classified in the mucilage metagenome to taxonomically classified diazotroph genomes validated our strategy to recover taxa with both high relative abundance in the aerial root mucilage microbiome and functionally important traits. Notably, the majority of genomes in our collection were classified to the Actinobacteria, Firmicutes, and Proteobacteria phyla, which strongly aligns with previous efforts to characterize plant-associated microbiomes (S1 Fig, S4 Table) (50-52). Reads classified to *Pseudmonas* in OLMM00 had the highest relative abundance among genera in the metagenome, and this isolate collection contained several distinct clades of *Pseudomonas* based on the substantial genome dissimilarity observed from all-by-all whole genome sequence comparisons (Fig 1). Whole genome taxonomic classification of diazotroph genomes also revealed presence of the second most abundant genus of OLMM00, *Acidovorax*, in the collection, as well as others assigned to genera with high relative abundance in the mucilage metagenome that include *Agrobacterium, Herbaspirillum* and *Burkholderia*. However, the majority of classified diazotrophs were Gammaproteobacteria that exhibited low relative abundance in OLMM00 (S1 Fig, S2 Fig, S7 Table). This suggested that diazotrophic contributions to Sierra Mixe maize by the mucilage microbiome may originate from community members of lower abundance, as evidenced by the diverse set of diazotrophic isolates described here. Furthermore, comparison of taxonomic analysis between whole genome sequences of selected diazotrophs and the OLMM00 metagenome suggested that microbial diversity of the mucilage microbiome is much broader than that of the collection. This suggests that diazotrophy may not be a widespread feature among genera detected in the OLMM00 mucilage metagenome.

### Diazotrophs exhibited the genomic potential for mucilage polysaccharide utilization

Utilizing the canonical pathway for BNF, one of the most energy-intensive biochemical processes in biology that consumes 16 ATP per reaction cycle to convert a single dinitrogen molecule into ammonia (53), an actively fixing diazotroph associated with Sierra Mixe maize would require a reliable feedstock to produce chemical energy. Based on the diverse monosaccharide composition (arabinose, fucose, galactose, glucuronate, mannose, xylose) of aerial root mucilage polysaccharide (6, 54) and evidence of endogenous GH activity present in fresh mucilage samples (55), we surmised that harnessing it for energy to drive BNF requires bacterial genes encoding both GHs to facilitate polysaccharide catabolism, and those conferring the ability to transport smaller sugars into the cell. We mined isolate genomes for carbohydrate utilization genes and parsed relevant data using manually curated lists of relevant database accessions (S9 Table and S11 Table) (33).

GHs are the most abundant class of carbohydrate active enzymes (CAZymes) and consist of over 150 distinct families with documented substrate specificities (56). Importantly, GHs often attribute multiple substrate specificities while maintaining similar protein domain architectures and sequence similarity. This ascribes the potential for substrate promiscuity among GH enzymes classified to a given GH family based on differences in protein structure. The GH profiles of isolate genomes indicated that mucilage diazotrophs possess the genomic potential to liberate monosaccharide components of the mucilage polysaccharide (Fig 2A). A summary of diazotrophic isolate counts for the number of isolates with genes in each GH group by genus classification further suggested that the majority of isolated diazotroph genomes encode highly specific as well as promiscuous GHs (S10 Table). These results indicated that mucilage diazotrophs are capable of liberating multiple polysaccharide derivatives irrespective of taxonomic assignment.

While the ability to liberate small carbohydrates from mucilage polysaccharide is necessary for its utilization as an energy source, diazotrophs from this niche must also possess the corresponding sugar transport systems. Bacteria possess multiple mechanisms for monosaccharide transport that primarily consist of membrane bound permeases, symporters, ABC-type porters and phosphotransferase (PTS) systems (57). We found the presence of sugar transporters from these classes with specificities for all six monosaccharide derivatives of mucilage polysaccharide in all of the genomes (Fig 2B, S12). Considering these findings along with observations that mucilage diazotrophs possessed highly promiscuous GHs corresponding to the mucilage composition, we surmised that mucilage bacteria are theoretically capable of utilizing their endogenous carbohydrate utilization genes to derive energy from mucilage carbohydrates. Broadly, this analysis confirmed that the majority of our diazotrophic isolates possess genes that may confer the ability to derive energy from mucilage polysaccharide and provides additional support for the hypothesis that diazotrophs of the mucilage microbiota utilize the polysaccharide to drive BNF.

### Diazotrophs formed three distinct nitrogen fixation groups based on genome analysis

Based on the isolation strategy to enrich for diazotrophic bacteria from the mucilage microbiome and the confirmed BNF phenotypes of diazotrophic isolates, we hypothesized that the diazotrophic genomes contain the minimum set of *nif* genes proposed by Dos Santos (7). Remarkably, the collection contained a mixture of diazotrophs that were categorized into three groups: the DSP group of diazotrophs fully adherent to the Dos Santos model for essential *nif* gene content, a smaller group of SDS diazotrophs with incomplete versions of the Dos Santos model, and the DSN group that completely lacked all six essential *nif* genes (Fig 3, S3 Fig, S4 Fig and S5 Fig). While the DSP group consisted of diazotrophs that possessed homologous sequences to HMMs for all six essential *nif* genes (*nif*HDKENB) of the Dos Santos model along with matches to the majority of other NIF regulon genes (7), discovery of the DSN and SDS isolates lacking homologous sequences to this set of canonical *nif* genes either entirely, or in-part, was unexpected. Interestingly, absence of matches to the HMM for the *nif*L gene that confers repression of the *nif*-specific transcriptional activator NifA in a large number of DSP diazotroph genomes suggests that these isolates may be acclimatized to high frequencies of nitrogen-fixing conditions in their native environment (58). Furthermore, the *nif*W gene was found to be non-essential for a large number of DSP diazotrophs that lacked presence of a homologous gene in their genome, which is corroborated by a previous report in *nif*W^-^ strains of *K. pneumoniae* (59). However, observations that all confirmed diazotrophs in the DSP group were adherent to the the well established genetic structure of the *K. pneumoniae* NIF regulon (44), and that genomes classified as *Klebsiella* were only assigned to the DSP group validated use of the *Klebsiella* model to examine the diazotrophic isolate genomes for canonical *nif* genes.

Taxonomic classification of diazotrophic genomes revealed a spectrum of phylogenetic diversity that was not found to be indicative of *nif* gene presence. For example, while gammaproteobacterial genera classified among DSP genomes included *Enterobacter, Klebsiella, Kosakonia, Metakosakonia, Pseudomonas, Rahnella* and *Raoultella*, the SDS and DSN groups contained genomes that were classified as *Enterobacter, Metakosakonia, Pseudomonas* and/or *Rahnella* as well. Our discovery of diazotrophs in the DSP group classified as Gammaproteobacteria suggested that bacteria of this taxonomic class from the mucilage environment are likely to contribute to the BNF phenotype of Sierra Mixe maize. This is supported by previous studies describing species from enterobacterial genera classified among genomes in the DSP group (*Enterobacter, Klebsiella, Kosakonia, Rahnella*, and *Raoultella*) as diazotrophic endophytes associated with cereal crops such as sugarcane, rice, and maize (60-64). Recent reports demonstrated the successful engineering of a *Pseudomonas* strain capable of associating with wheat and maize as a diazotrophic endophyte (65), as well as successful growth promotion of maize using a diazotrophic strain of *Pseudomonas* isolated from the rhizosphere of rice (66). However, to the best of our knowledge, a naturally occurring diazotrophic pseudomonad associated with maize endophytically is yet to be reported. Additionally, genomes in the SDS and DSN NIF groups were classified to many other genera outside of Gammaproteobacteria, which indicates that diazotrophs of Sierra Mixe maize exhibit much broader phylogenetic diversity relative to these previous reports of diazotrophs that associate with cereal crops.

### Many diazotrophs exhibited high BNF ratios independent of possessing canonical *nif* genes

In contrast to our hypothesis, results from the BNF assay and *nif* gene mining confirmed a substantial portion of the isoated diazotrophs lacked homologous protein coding sequences to many, or all, canonical *nif* genes of the Dos Santos and *Klebsiella* models yet exhibited high BNF ratios independent of canonical *nif* genes. Our quantitative assay to detect the incorporation of ^15^N-dinitrogen from an enriched atmosphere into secreted metabolites served as a robust alternative to conventional methods of diazotrophic detection, such as colorimetric assays for ammonium secretion and the acetylene reduction assay, which limit detection of evidence for BNF to ammonium accumulation or secondary nitrogenase activity (i.e. production of ethylene through the reduction of acetylene gas), respectively (67, 68). As there has never been a documented case of diazotrophs utilizing atmospheric nitrogen without key components of the nitrogenase enzyme complex, our observations that SDS and DSN diazotrophs lacked protein coding sequences homologous to essential *nif* genes in their genomes (S4 Fig, S5 Fig) lead us to question the metabolic mechanisms that allowed them to be successfully cultured and isolated on nitrogen-free medium in the laboratory.

While comparison of *nif* gene profiles (Fig 3) with results from the BNF assay confirmed that DSP isolates utilize atmospheric nitrogen for growth, comparison with BNF assay results for the SDS and DSN NIF groups indicated that these isolates were also capable of incorporating atmospheric nitrogen into secreted metabolites at efficiencies that both rivaled and exceeded those of DSP isolates in some cases (Fig 4). For example, while lactococci are commonly associated with plants (69), our investigation serves as the first report of diazotrophic lactococci based on observations that *Lactococcus* isolates exhibited some of the highest BNF ratios (Fig 4C, S2 Table). These results were unexpected due to the total absence of homologous sequences to HMMS for essential *nif* genes within lactococcal isolate genomes (Fig 3, S5 Fig), and suggested that bacteria of the mucilage microbiota lacking essential *nif* genes are capable of incorporating atmospheric nitrogen into their metabolism under N-limiting environmental conditions through metabolic mechanisms outside of the Dos Santos and *Klebsiella* models. Taken together, the genome analysis and BNF assay results revealed that possession of canonical *nif* genes comprising the Dos Santos and *Klebsiella* models were not required for all diazotrophs from Sierra Mixe maize to exhibit BNF activity, suggesting that novel diazotrophic mechanisms exist in this community.

Uncovering the genetic underpinnings of the observed BNF phenotype for mucilage diazotrophs lacking canonical *nif* genes will rely on advances in genomic analysis and future experimentation. While HMMs derived from consensus sequences of full-length coding sequences serve as a reliable tool to detect known genomic features in bacteria, they do not invite the possibility of detecting novel protein coding sequences conferring known biological functions through alternative protein domain architecture. Therefore, advances in genome annotation that integrate machine learning algorithms with HMM libraries derived from consensus sequences of protein domains rather than full-length coding sequences, such as *Nanotext*, may enable the discovery of new proteins conferring familiar activities (70). Additionally, implementation of microbial pan-genome association studies using appropriate control groups for DSN isolates with confirmed BNF phenotypes may also shed light on additional significant genes associated with diazotrophy (71).

### Alternative nitrogenase genes were not present in SDS and DSN isolate genomes

We queried WGS from diazotrophic isolates for protein coding sequences homologous to known alternative nitrogenase genes in search of an explanation for the discovery that confirmed diazotrophic isolates lacked essential *nif* genes of the Dos Santos and *Klebsiella* models. Environments with limited abundance of molybdenum often harbor diazotrophic bacteria that exhibit genetic operons encoding alternative nitrogenase systems. These include Vanadium-Iron (Vn-Fe) type and Iron-only type nitrogenases (Fe-Fe) that assume quaternary structure without utilization of molybdenum and the assistance of an additional *nif* gene encoding the *gamma* subunit for the catalytic component (43). Additionally, these operons arose over evolutionary time through genetic duplication events and neofunctionalization of the Fe-Mo *nif*HDK operon in response to abiotic stress (43, 53). Referencing previous reports on the nutrient deficient quality of indigenous fields for Sierra Mixe maize cultivation (6), the BNF assay, and *nif* gene mining results, we hypothesized that SDS and DSN diazotrophs possessed alternative *nif* genes and tested it by scanning the protein coding sequences of diazotroph genomes with HMMs for the Vn-Fe *nif* genes (*vnf)* and Fe-Fe *nif* genes (*anf*).

While results from this investigation forced the rejection of our hypothesis by confirming that SDS and DSN isolates do not possess alternative *nif* genes, they did reveal discovery of a subset of diazotrophs from the DSP group that possessed genes resembling the *anf* and *vnf* genetic operons. We found 42 diazotrophs with genes matching TIGRFAMs from all three classes of known nitrogenase systems (Fig 5A). Although unexpected, this result corroborates the previous report that alternative *nif* genes were only found to occur in diazotrophs that also possessed the Mo-Fe nitrogenase system (53), and the observation of alternative *nif* gene sequences in Sierra Mixe mucilage (6).

Comparison of whole genome nucleotide composition for diazotrophs with homologs to alternative *nif* genes provided evidence that this subset of the DSP NIF group exhibited considerable genomic diversity and contained distinct members with resemblance to previously reported *Metakosakonia* and *Raoultella* reference genomes (Fig 5B). However, this subset of diazotrophic isolates exhibited high genome dissimilarity and the group was found to contain genomes for which assignment to a known genus was unattainable through LCA classification using the GTDB. These observations suggested that the mucilage microbiota harbors *Metakosakonia* and *Raoultella* with alternative *nif* genes and variation in metabolic capabilities, as well as potentially novel genera with considerable genomic differences. Further investigation by pan-genome analysis revealed large blocks of genomic features corresponding to the variation in genome composition observed in four isolate genomes that formed a distinct clade (Fig 5C). To our knowledge, this is the first report of maize-associated *Raoultella* exhibiting alternative *nif* genes, and the genomic evidence surrounding this discovery invites the possibility for classification of a new species within the genus.

This work reaffirmed the proposal of Sierra Mixe maize as a model system to investigate nitrogen fixation in cereal crops by validating its association with diazotrophic bacteria that possess canonical genetic operons for nitrogen fixation (72). Our investigation emphasized the importance of aerial root mucilage to the nitrogen-fixing phenotype of the system by confirming the presence of classical nitrogen fixing bacteria in the aerial root mucilage microbiota that contained the genomic potential to derive energy for BNF from mucilage polysaccharide. We also demonstrated that mucilage-derived diazotrophs incorporated atmospheric nitrogen into their metabolism through unknown metabolic pathways extending beyond current knowledge that defines BNF as bacterial conversion of dinitrogen to ammonia through the expression of canonical *nif* gene products within the Dos Santos and *Klebsiella* models. We succeeded in recovering and characterizing diazotrophs from the mucilage microbiota and found diazotrophs that did not contain any canonical *nif* genes, suggesting their use of novel genes for the conversion of dinitrogen into organic nitrogen forms that were assimilated into many small molecules exported by the organisms. Collectively, this study demonstrated that specific microbiome members of Sierra Mixe maize display diazotrophy with multiple molecular mechanisms.

## Funding information

Mars, Incorporated http://www.mars.com/global. The research was funded by an unrestricted gift and a grant to ABB and BCW. The funder had no role in study design, data collection and analysis, decision to publish, or preparation of the manuscript. The research was funded by grants to ABB from BioN2, Incorporated. The funder had no role in study design, data collection and analysis, decision to publish, or preparation of the manuscript. The research was also funded by United States Department of Agriculture (USDA) by a grant to BCW (award #2019-67013-29724).

## Author Contributions

BCW and MY and NK carried out the strategy to culture, isolate and store the microbial collection from Sierra Mixe maize. SMH, TP and BH constructed DNA sequencing libraries for WGS of bacterial isolates and BH conducted the genome sequencing. SMH carried out all bioinformatic analyses related to WGS analysis with guidance from BCW and CTB. BCW and RJ designed the BNF assay, established the method and analyzed the associated data, and NK conducted the experiments. SMH wrote the first draft of the manuscript. ABB, BCW, CTB, SMH and TP edited and revised the manuscript.

## Conflict of interest

The authors declare no conflicts of interests. None of the authors are employed by the major funding agency of this work, MARS, Inc.

## Notes

### Competing Interest Statement

The authors have declared no competing interest.

