## supporting information for "Genomic characterization of a diazotrophic microbiota associated with maize aerial root mucilage"

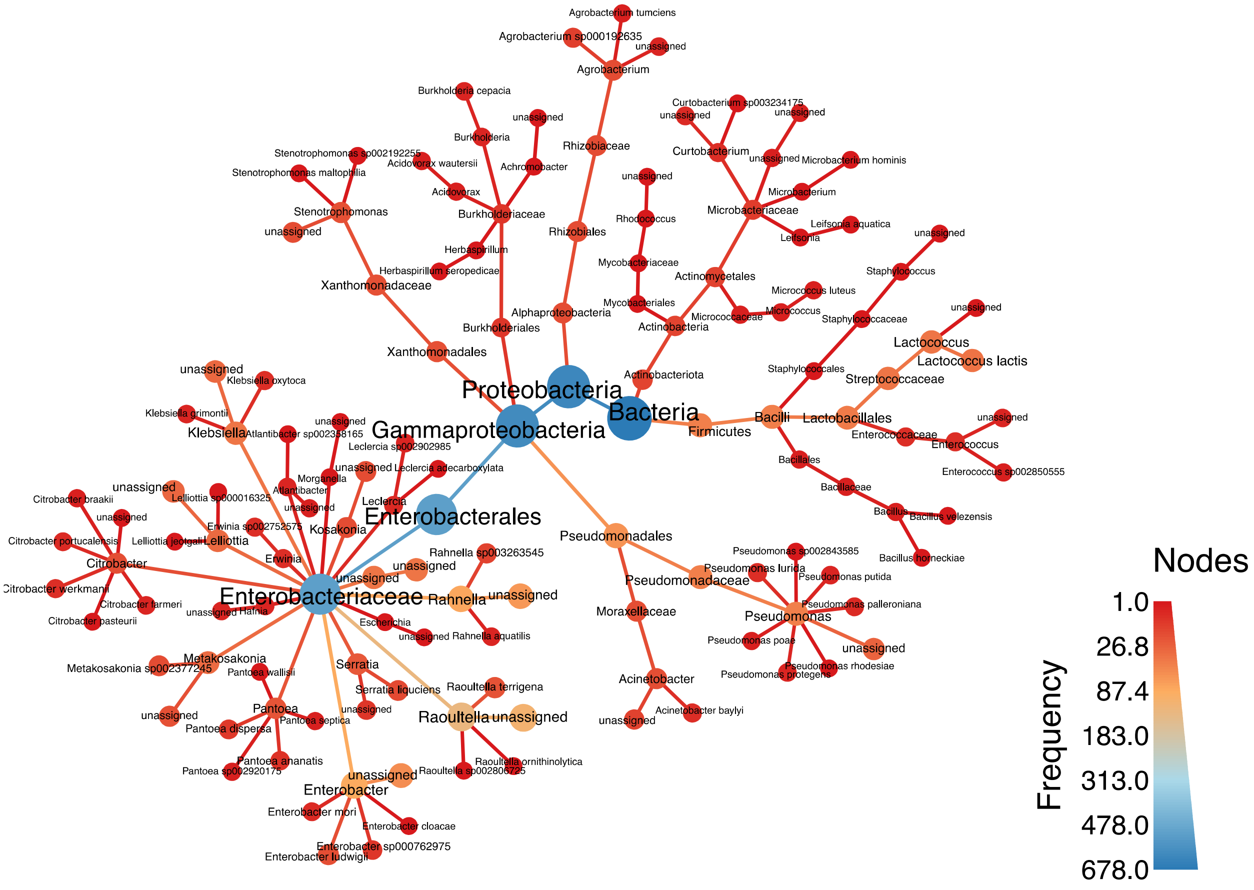


**S1 Fig. Taxonomic classification of genome bins from mucilage isolates.** Heat Tree showcasing the phylogenetic diversity of all Metabat (1) genome bins derived from isolate draft genome assemblies. Node size corresponds to the frequency of occurrence at each level of taxonomic classification. Dark blue corresponds to the highest observation count of a taxonomic assignment and dark red indicates low observation frequency.

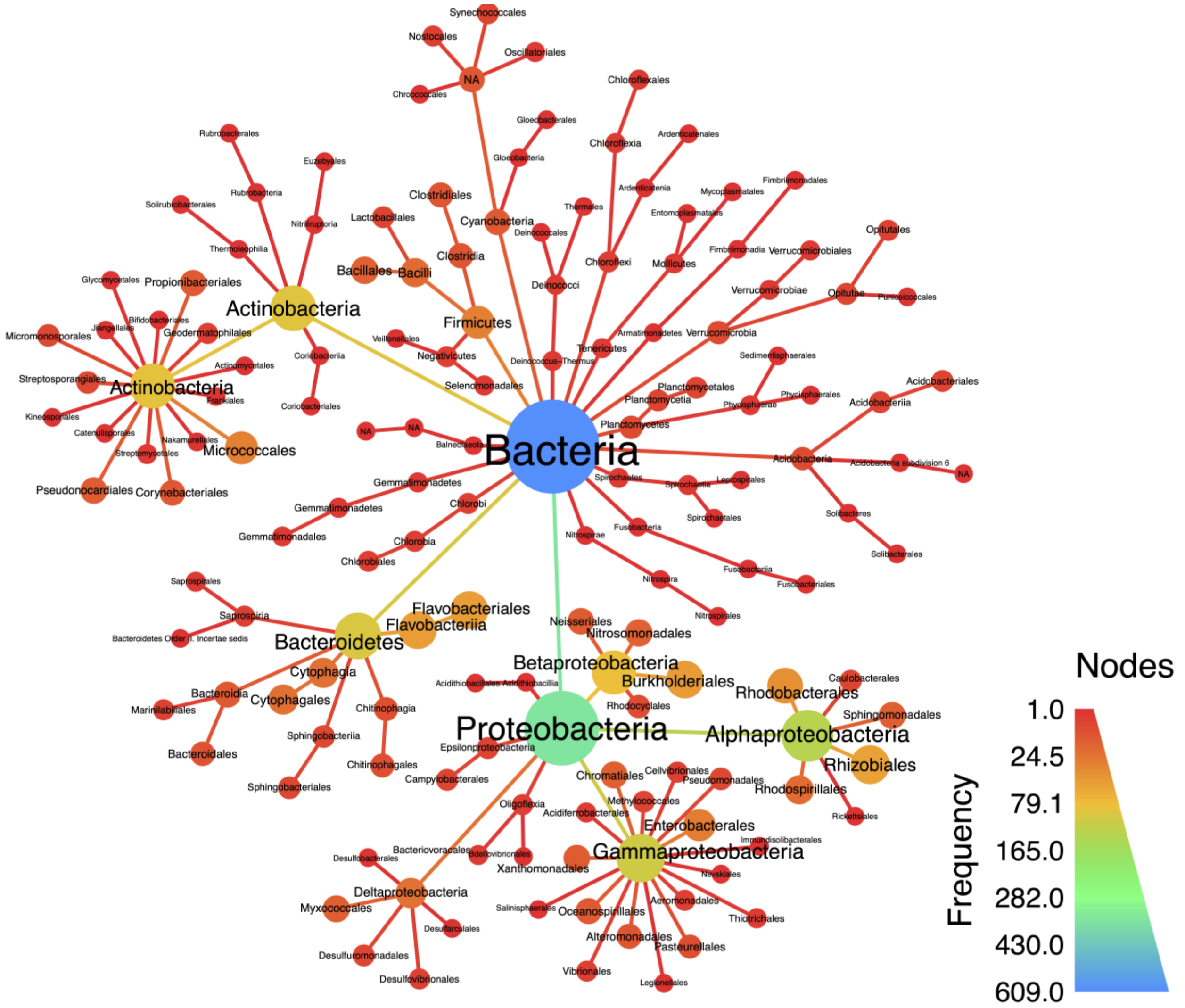


**S2 Fig. Observation counts of classified taxa in the mucilage metagenome.** Quality trimmed short reads of the OLMM00 mucilage metagenome were input to Kraken2 (2) and classified using the RefSeq complete database for microbial genomes installed with built-in commands of Kraken2. Bracken2 (3) was used to re-estimate classified read counts and the data was imported to R in biom-format. Taxa with readcounts less than 500 were filtered using Phyloseq (4) and the data were visualized using MetacodeR (5). Terminal nodes represent taxa classified at the order level and node size corresponds to frequency of observation for each taxonomic level.

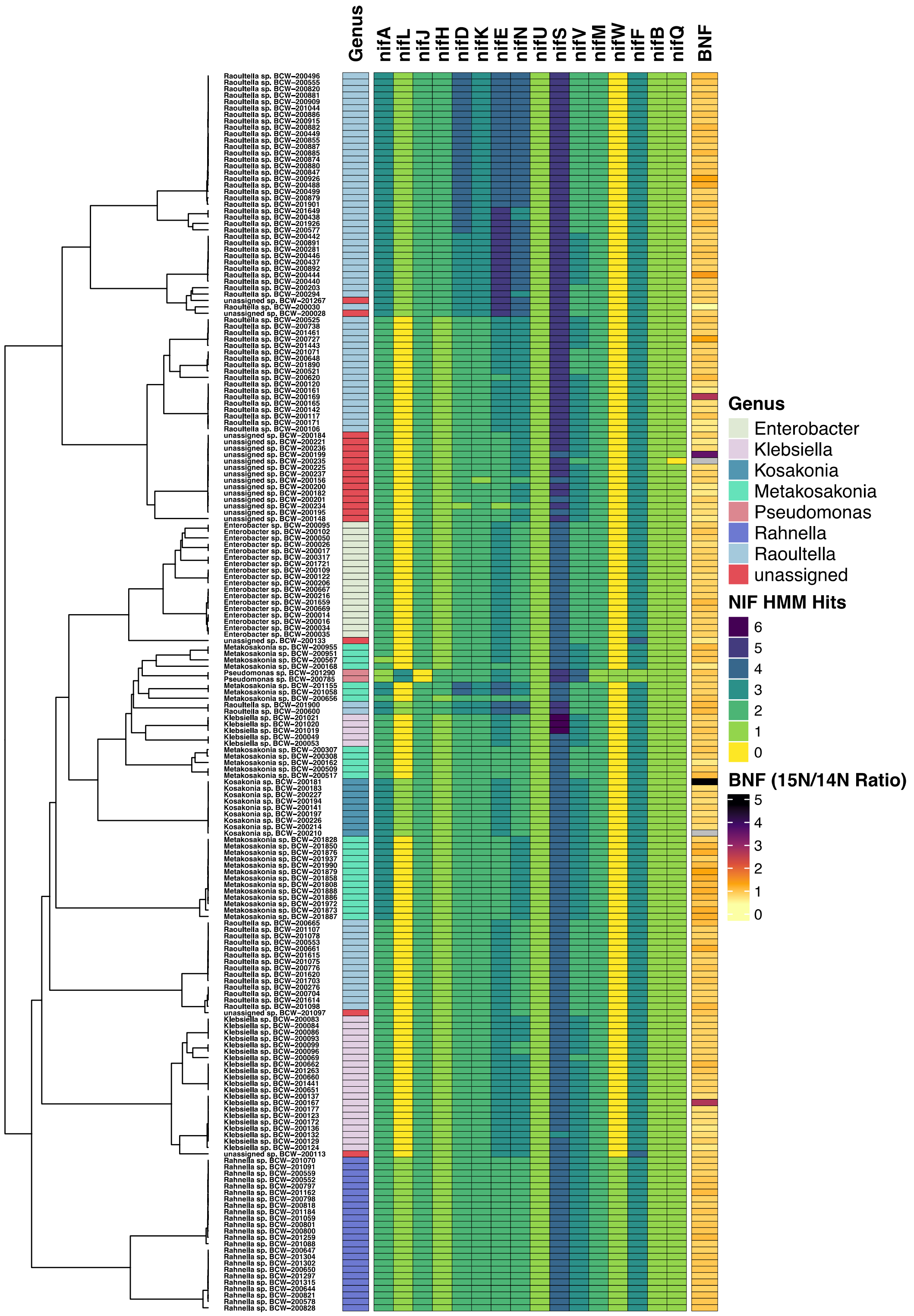


**S3 Fig. *Nif* gene profile and BNF assay performance of Dos Santos Positive isolates.** *Nif* gene profiles for isolate genomes of the DSP group were extracted and visualized independently using the ComplexHeatmaps (6) package in R. Isolate genomes were clustered using the dendrogram output from Sourmash (7) and each genome row is presented along with annotations to indicate lowest common ancestor (LCA) classification data at the genus level.

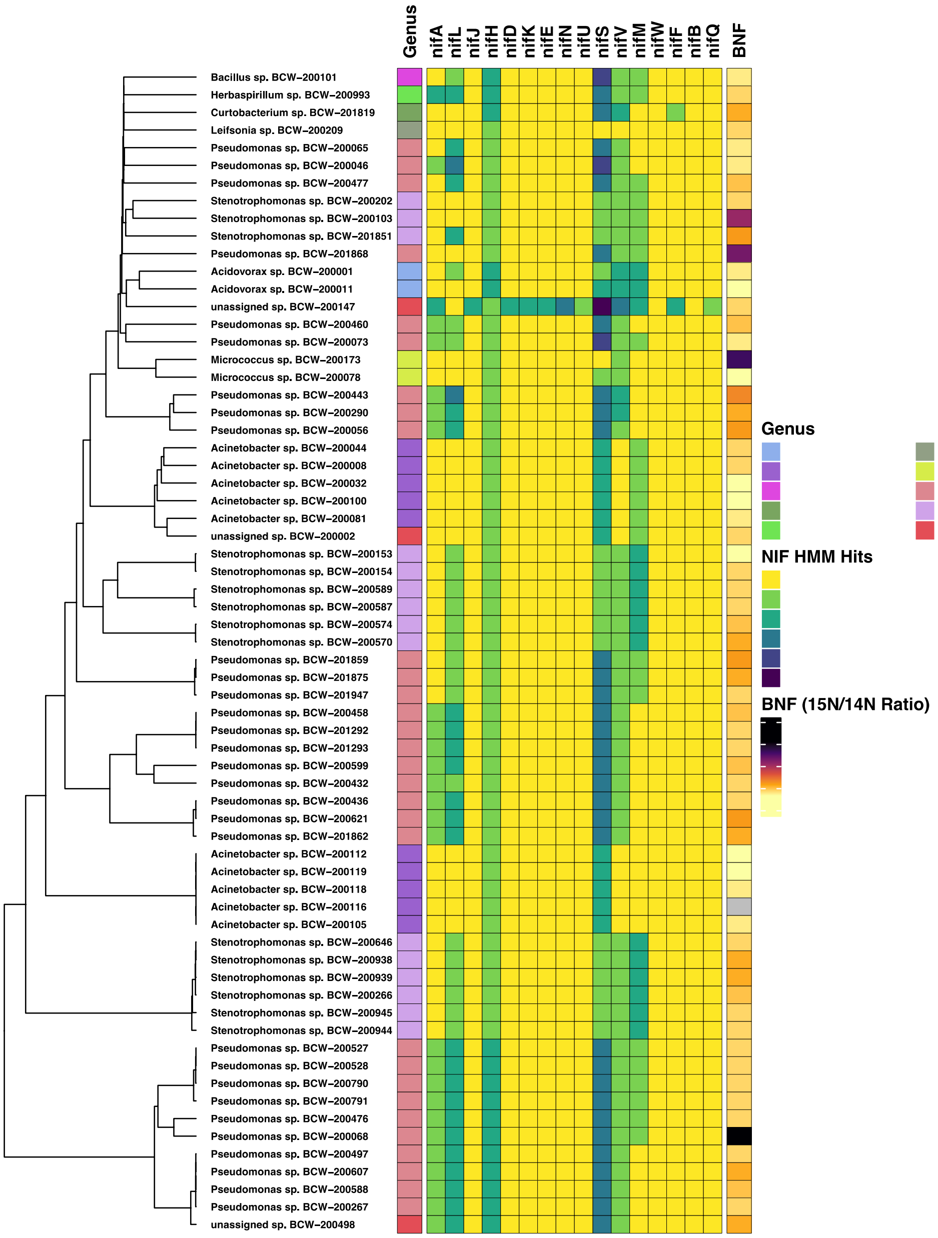


**S4 Fig. Nif gene profile and BNF assay performance of Semi-Dos Santos isolates.** *Nif* gene profiles for isolate genomes of the SDS group were extracted and visualized independently using the ComplexHeatmaps (6) package in R. Isolate genomes were clustered using the dendrogram output from Sourmash (7) and each genome row is presented along with annotations to indicate LCA classification data at the genus level.

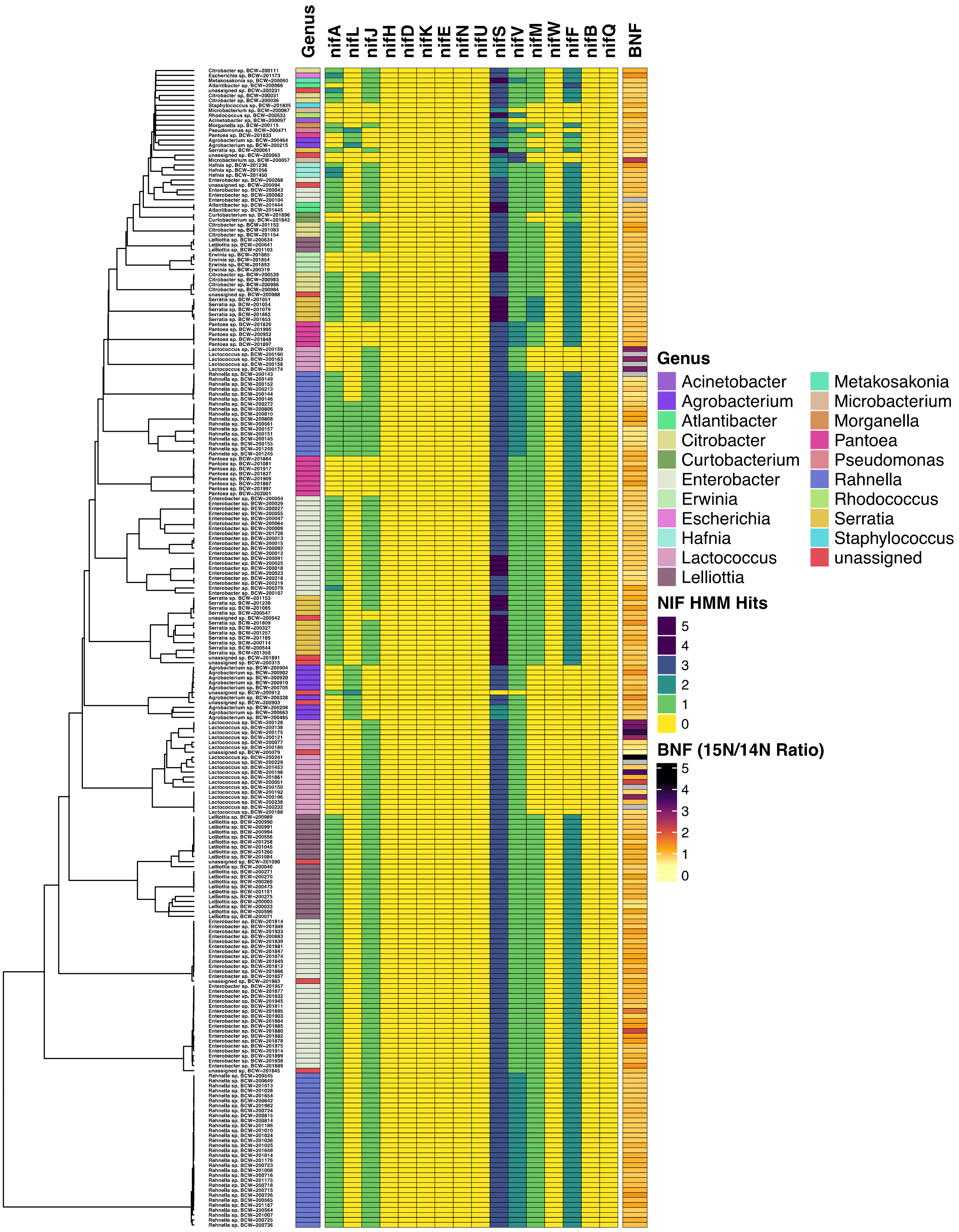


**S5 Fig. Nif gene profile and BNF assay performance of Dos Santos Negative isolates.** *Nif* gene profiles for isolate genomes of the DSN group were extracted and visualized independently using the ComplexHeatmaps (6) package in R. Isolate genomes were clustered using the dendrogram output from Sourmash (7) and each genome row is presented along with annotations to indicate LCA classification data at the genus level.

### Supplementary Tables

Supplementary tables are provided as sheets within a single Microsoft excel workbook. Sheet names correspond to the following table legends that provide additional information related to the values presented in each table.

### Supplementary Table Legends

**S1 Table. Culturing conditions for Sierra Mixe maize bacterial isolates.**

Isolation sources included samples collected from landrace maize varieties grown in the Sierra Mixe region of Oaxaca, Mexico. Medium types included Blood-Heart-Infusion Agar (BHI), M9 minimal medium, and a custom nitrogen-free minimal medium (NFM).

**S2 Table. List of diazotrophic isolates and their BNF ratios.** $^{15}N/^{14}N$ ratios (BNF ratios) for each isolate are presented alongside their unique isolate ID number. BNF ratios represent the quotient of summations for peak intensities of all statistically significant N-containing biomarkers under both enriched and non-enriched atmospheric conditions. N-containing biomarkers were determined by analysis of LC-MS data using Metaboanalyst (8). See methods for further details on determination of N-containing biomarkers.

**S3 Table. Bacterial whole genome sequencing library and assembly metrics.** Genome assemblies generated with MEGAhit (9) were assessed using QUAST (10) to generate the summary data presented that include number of contigs, total length of the assembly in megabase pairs (Mb), length of the largest contig in the genome assembly in kilobase pairs (Kb), percentage of the genome comprised by guanine and cytosine (GC %). Mean fold coverage was computed by mapping input reads back to the genome assembly and computing an average value for the coverage of all contigs in the assembly. Genome bins were generated using Metabat (1).

**S4 Table. Taxonomic classification of all sequenced bacterial isolate genomes.** All sequenced isolate genomes were classified using the 'lca classify' function of Sourmash (7). MinHash sketches were generated using a k-size of 31 and a scaled value set at 2000. Each genome sketch was queried against a database of MinHash sketches for Genome Taxonomy Database (GTDB) (11) version 89 (available at: https://osf.io/gs29b/).

**S5 Table. Taxonomic classification of WGS bacterial isolate bins.** Genome bins from all sequenced isolate genomes were generated using Metabat. These bins were subsequently classified using the 'lca classify' function of Sourmash (7). MinHash sketches were generated using a k-size of 31 and a scaled value set at 2000. Each genome sketch was queried against a database of MinHash sketches for Genome Taxonomy Database (GTDB) (11) version 89 (available at: https://osf.io/gs29b/).

**S6 Table. Summary of pure isolate WGS.** Pure isolate genome assemblies were generated using MEGAhit (9), assessed using QUAST (10), and taxonomically classified using the 'lca classify' function of Sourmash (7) (see Methods). MinHash sketches were generated using a k-size of 31 and a scaled value set at 2000. NIF Group assignments indicate that isolate genomes were either positive for the presence of all six essential *nif* genes of the Dos Santos Model (DSP), had a semi-complete set of the six *nif* genes (SDS), or were negative for all six (DSN) (12).

**S7 Table. Mucilage metagenome taxonomic classification and normalized readcounts.** The OLMM00 mucilage metagenome previously reported by Van Deynze et al. in 2018 (13) was re-analyzed using Kraken2 (2) and the RefSeq complete database of microbial genomes (14). Classified read counts were re-estimated using Bracken2 and exported in biom-format using Kraken-biom. The read counts were imported to R and normalized to counts per million using Phyloseq. Taxa with fewer than 500 classified reads were removed. Relative abundances were generated by dividing re-estimated read counts for each taxon by the sum total of all classified reads post-filter.

**S8 Table. Summary of mucilage metagenome taxonomic observation frequencies.** This table provides the corresponding values for taxonomic observation frequencies in the OLMM00 mucilage metagenome after re-estimation of Kraken2 (2) classified read counts by Bracken2 (3) that were used to generate S1 Fig.

**S9 Table. Carbohydrate Active Enzyme Families used for CAZyme genome screening.** The CAZy database (15) was referenced to generate a manually curated list of CAZyme families that were determined to by relevant for utilization of mucilage polysaccharide. Reported substrate specificities were considered for each family to form eleven custom groupings based on sugar residue activities.

**S10 Table. Summary of GH family gene presence in pure isolate genomes.** Values indicate the number of isolates with one or more unique genes identified within the genome that matched a HMM for GH families within a designated grouping: Ara – GH(62,127,137,142,146); Ara/Gal/GlcA/Man/Xyl - GH2; Ara/Xyl - GH(3,43,51,54); Fuc - GH(29,95,139,141,151); Fuc/GlcA/Xyl - GH30; Gal - GH(16,27,35,36,42,57,59,95,97,98,110,147,160,165); Gal/GlcA - GH4; Gal/GlcA/Fuc/Man/Xyl - GH1; Gal/Man/Xyl - GH 31; GlcA - GH(67,79,115,154); Man - GH(5,38,47,63,92,99,125,130,164). 'N isolates' indicates the total number of pure isolate genomes that were classified to the indicated genus and included in the analysis.

**S11 Table. TCDB Accessions used for sugar transport gene detection.** Transporter accession IDs, transporter type and descriptions were reported based on information available in the Transporter Classification Database (www.tcdb.org) (16). Sugar transporters were selected by manually searching through the TCDB for bacterial sugar transport genes that corresponded to monosaccharide components of the mucilage polysaccharide from Sierra Mixe maize.

**S12 Table. Summary of sugar transporter gene presence in pure isolate genomes.** Values represent the total number of isolates with detected membrane transporters for each sugar component of mucilage polysaccharide. 'N Isolates' corresponds to the number of isolate genomes sequenced under the given taxonomic assignment at the genus level. Transporter mechanisms for each sugar were grouped based on the type of sugar transport to calculate sum totals for each isolate class.

**S13 Table. Summary for the Pan-genome of isolates possessing alternative *nif* genes.** The pan-genome for isolates containing alternative *nif* genes was generated using Roary 3.12.0 (17). The summary of features was generated as standard output from running the bioinformatic pipeline with GFF files for each isolate generated using Prokka 1.12 (18).
